# The ALS- and FTD-associated proteins Annexin A11 and CHMP2B act sequentially in membrane repair

**DOI:** 10.1101/2024.11.19.624330

**Authors:** C.M. Heffner, G.P. Starling, A.M. Isaacs, J.G. Carlton

**Affiliations:** Organelle Dynamics Laboratory, The Francis Crick Institute, 1 Midland Road, London, NW1 1AT; School of Cancer & Pharmaceutical Sciences, King’s College London, London, SE1 1UL; UK Dementia Research Institute at UCL, London, WC1E 6BT; Department of Neurodegenerative Disease, UCL Queen Square Institute of Neurology, London, WC1N 3BG

## Abstract

Maintenance of plasma membrane and organellar integrity is essential for cell viability. Cells must recognise damaged membranes and orchestrate repair programmes to preserve compartmentalisation. A variety of cellular factors including ESCRTs, Annexins, stress granules, lipids and proteins allowing vesicle and organelle fusion with damaged membranes have been reported to contribute to membrane repair. However, whether these factors operate independently or together to repair membranes is unclear. Here, we expose temporal differences and interdependencies in the recruitment of ESCRT-III and Annexin proteins to sites of membrane damage. We show that while Annexins are recruited immediately to sites of damage, ESCRT-III assembles only after membrane sealing. We show that ESCRT-III acts to shed damaged membranes from the cell and that FTD-and ALS-associated mutations in CHMP2B and ANXA11 compromise the repair process. These data present an integrated ‘sealing and healing’ model of events allowing membrane repair and restoration of membrane integrity.

**One-Sentence Summary:** A rubric of sealing and healing for ESCRT-mediated membrane repair

## Introduction

Maintenance of plasma membrane and organellar integrity is essential for cellular viability, the maintenance of ionic gradients across membranes and the compartmentalisation of proteins and bioactive molecules. Membranes can be damaged physically by the forces imposed by cell and tissue mechanics, crystalline material or aggregates, including those associated with neurodegenerative disease^1,2^ such as amyloid-beta^3,4^, alpha synuclein^5^ and tau-fibrils^6^ and the activities of pathogens such as *Salmonella* or *Mycobacterium*^7^. Membranes can be damaged chemically through reactive oxygen species^8,9^, and biochemically, through the actions of bacterially secreted Pore Forming Toxins (PFTs), cellular components of immune control such as Perforins^10^, Gasdermins^11–15^, Ninjurin-1^16,17^ and the complement membrane attack complex^18^ . Reactive oxygen species^8,9^, are also thought to damage cellular membranes. A variety of mechanisms to repair membrane damage have been proposed, including the Endosomal Sorting Complex Required for Transport-III (ESCRT-III) machinery, Annexins, delivery of lipids by membrane contact sites, vesicular or organellar fusion and the assembly of stress granules at sites of damaged membranes^19,20^. Whether these activities represent truly independent membrane repair mechanisms or are integrated parts of a singular process is currently unclear. ESCRT-III is an ancient membrane remodeling complex with essential roles in several membrane shaping processes throughout the cell including endolysosomal sorting, exosome biogenesis, enveloped retrovirus release, cytokinetic abscission, nuclear envelope regeneration during mitotic exit and the repair of ruptured nuclear, plasma and lysosomal membranes^21–26^. Mammalian cells express 11 different ESCRT-III subunits called Charged Multivesicular Body Proteins (CHMPs) and a related protein called IST1, that, alongside an AAA-ATPase called VPS4, act together to orchestrate the membrane remodeling process^27^. C-terminal truncating mutations in the ESCRT-III subunit CHMP2B (CHMP2B^INT5^ or CHMP2B^Q165X^) are rare causative factors of autosomal dominant Frontotemporal Dementia (FTD)^28–30^. How these mutations lead to disease is currently unknown. Several other ESCRT-associated proteins have been implicated in a range of neurodegenerative diseases and neurodevelopmental disorders, suggesting a wider implication of ESCRT activity in neurological health^31–35^. The calcium dependent phospholipid binding Annexin proteins have also been implicated in membrane repair at plasma, endosomal and nuclear membranes^36–39^. Mutations in Annexin A11 (ANXA11) are associated with Amyotrophic Lateral Sclerosis (ALS)^40^, a disease with genetic, clinical and neuropathological overlap with FTD^41^. Given the involvement of lipid regulation in neurodegeneration^42^ and recent suggestions that membrane damage is at the core of Alzheimer’s disease^43^, we wondered if these different FTD/ALS risk factors operated in a common pathway to provide protective functions that preserve cellular viability. We used advanced live cell imaging to examine CHMP2B^INT5^ and CHMP2B^Q165X^ in response to multiple forms of plasma membrane damage and describe previously unreported differences in their subcellular localisation that lead to temporal defects in their response. By imaging at high temporal resolution, we demonstrate sequential recruitment of Annexins and ESCRT-III to sites of membrane damage, with ESCRT-III arriving after membrane sealing has occurred. Further, we show that FTD-associated and ALS-associated mutations in *CHMP2B* and cause a phenotypically similar stalling of ESCRT-III at sites of membrane damage. These data provide an integrated ‘sealing and healing’ model for the repair of damaged cellular membranes to allow preservation of membrane integrity, and characterise a common mechanistic phenotype associated with separate FTD/ALS-associated mutations.

## FTD-causing mutations in CHMP2B cause its nuclear retention

To interrogate the cellular biology of the FTD-causing C-truncating mutations of CHMP2B, we first generated CAL-51 cells stably expressing low levels of GFP-tagged versions of human CHMP2 proteins (Figure 1A). We used the linker from the C-terminal Localisation and Purification (LAP) tag^44^, referred to as ‘L’, to preserve CHMP function. We were surprised to find that whilst CHMP2B-L-GFP was largely cytosolic, CHMP2B^INT5^-L-GFP and CHMP2B^Q165X^-L-GFP were both retained in the nucleus (Figure 1B and 1C). We confirmed these distributions using a C-terminal Hemagglutinin (HA) tag (Figure S1A and S1B). Extending these observations, we found that the CHMP2B paralog, CHMP2A-L-GFP, displayed a predominantly nuclear distribution (Figure 1C). These data show that the steady-state distribution of CHMP2A and CHMP2B between nucleus and cytoplasm is non-equivalent and that FTD-causing mutations in CHMP2B lead to its inappropriate nuclear retention.

**Figure 1.**
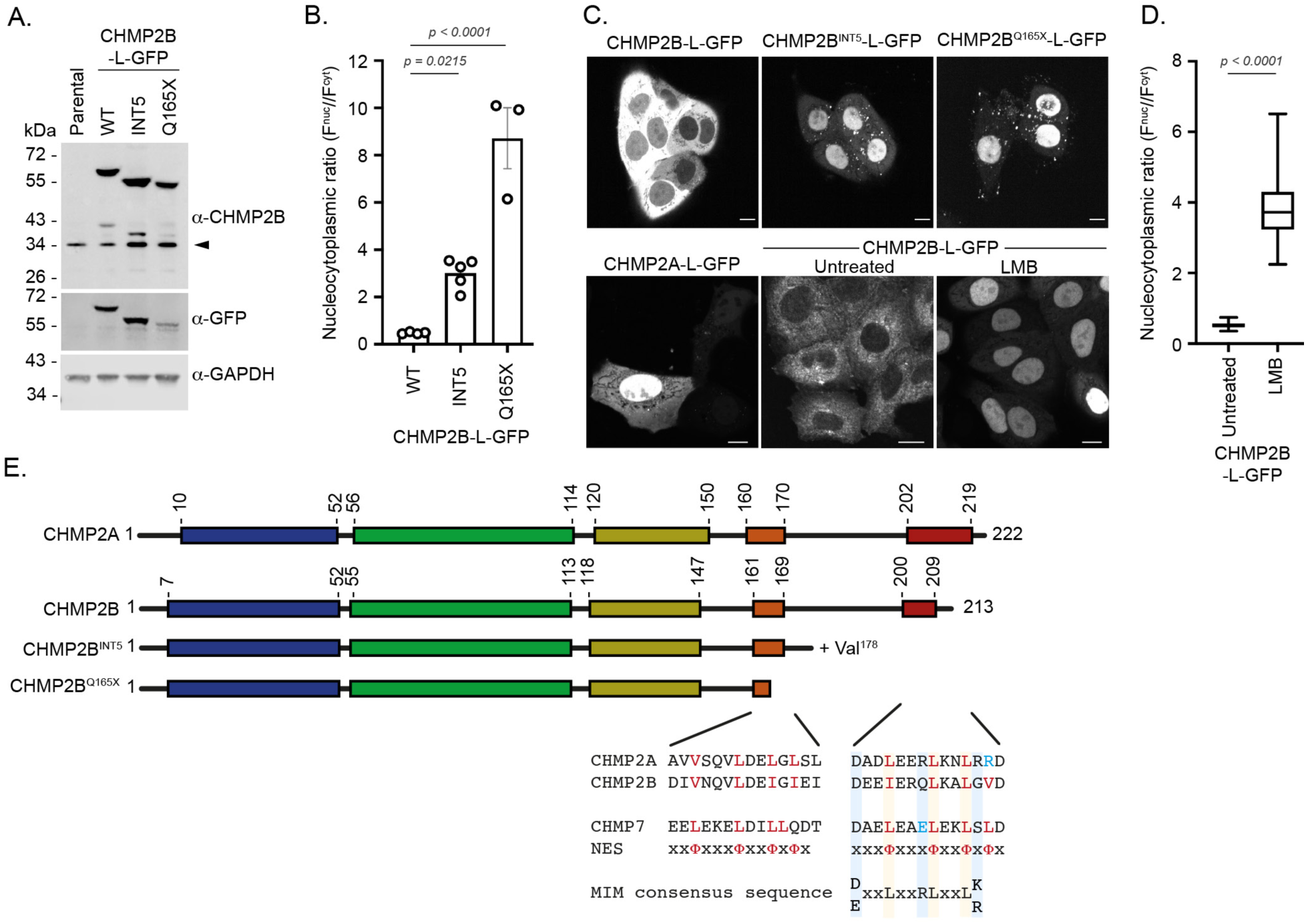
C-truncating pathogenic variants of CHMP2B are redistributed to the nucleus. **A**. Resolved cell lysates of CAL-51 cells stably expressing CHMP2B-L-GFP, CHMP2B^INT5^-L-GFP or CHMP2B^Q165X^-L-GFP were examined by western blotting with antisera raised against CHMP2B, GFP or GAPDH. Arrowhead indicates endogenous CHMP2B. **B**. Quantification of nucleocytoplasmic fluorescence from cells stably expressing CHMP2B-L-GFP (n = 57), CHMP2B^INT5^-L-GFP (n = 31), CHMP2B^Q165X^-L-GFP (n = 105). Mean ± S.E.M displayed from 3-5 independent experiments with statistical significance calculated by one-way ANOVA. **C**. Live cell imaging of CAL-51 cells stably expressing CHMP2B-L-GFP, CHMP2B^INT5^-L-GFP, CHMP2B^Q165X^-L-GFP or CHMP2A-L-GFP and imaging of CAL-51 cells stably expressing CHMP2B-L-GFP and treated with Leptomycin B (10 ng/mL, 4 hours). The nucleocytoplasmic ratio of CHMP2A-L-GFP (mean ± S.E.M. from 32 cells across N = 3 independent experiments) was 2.99 ± 0.71. Scale bar is 10 μm. **D**. Quantification of the nucleocytoplasmic ratio of GFP fluorescence in images from C (CHMP2B-L-GFP (n = 47 (untreated) or 101 (+ LMB) cells).. Data displayed as a box and whisker plot with median, 25^th^ and 75^th^ percentile displayed. Significance calculated by 2-tailed t-test. **E**. Schematic representation of CHMP2A, CHMP2B, CHMP2B^INT5^ and CHMP2B^Q165X^. Coloured blocks represent helix boundaries obtained from Alphafold2 predictions of CHMP2A (O43633) or CHMP2B (Q9UQN3) sequences. Key hydrophobic residues in Nuclear Export Sequences (NES) in CHMP2B’s C-terminal helices highlighted in red text, alongside equivalent residues from the CHMP7 NESs. Yellow shading used to indicate NES conservation, with the hydrophobic to basic amino acid changes in CHMP2A’s Helix 6 NES highlighted in blue text. Blue shading used to highlight key MIT-domain Interaction Motif (MIM) residues in Helix 6 of CHMP2A and CHMP2B, with the basic-to-acidic amino acid change of CHMP7’s MIM^45^ highlighted in blue.

Partitioning between nucleus and cytoplasm is governed by access to the nucleocytoplasmic transport machinery. We demonstrated that exportin inhibition with Leptomycin B (LMB) resulted in strong nuclear accumulation of CHMP2B-L-GFP (Figure 1C and 1D), but had limited effect on the nuclear retention of CHMP2B^INT5^-L-GFP and CHMP2B^Q165X^-L-GFP (Figure S1C - S1E). As such, we conclude that C-truncating mutations in CHMP2B restrict its nuclear export. Type-1 Nuclear Export Sequences (NES) in Helix5 and Helix6 of CHMP7 are critical to its control of the timely regeneration of the nuclear envelope during mitotic exit^45–47^. We noted two similar, but previously uncharacterised, Type-1 NESs in the C-terminus of CHMP2B that were truncated in CHMP2B^INT5^ and CHMP2B^Q165X^ (Figure 1E). By individually inactivating these sequences, we found that whilst both NESs contributed transport activity, the C-terminal NES was stronger than the internal NES (Figure S1F and S1G). This is consistent with a stronger nuclear retention of CHMP2B^Q165X^-L-GFP relative to CHMP2B^INT5^-L-GFP (Figure 2B and 2C) as it is missing both NESs. CHMP2A encodes a basic residue instead of a hydrophobic residue at position 221 and we hypothesised that this would inactivate its C-terminal NES, leading to its nuclear retention (Figure 1E). We mutated the equivalent residue (Val212) in CHMP2B and found that CHMP2B^V212K^-L-GFP was now retained in the nucleus (Figure S1F and S1G). Finally, we fused the Helix6 NES from CHMP7 to the C-terminus of CHMP2B^INT5^-L-GFP or CHMP2B^Q165X^-L-GFP (Figure S1H) and recovered cytoplasmic localization of these C-truncated versions of CHMP2B (Figure S1F and S1G). These data identify NES sequences in CHMP2B’s C-terminal helices that act to position this protein in the cytoplasm.

**Figure 2.**
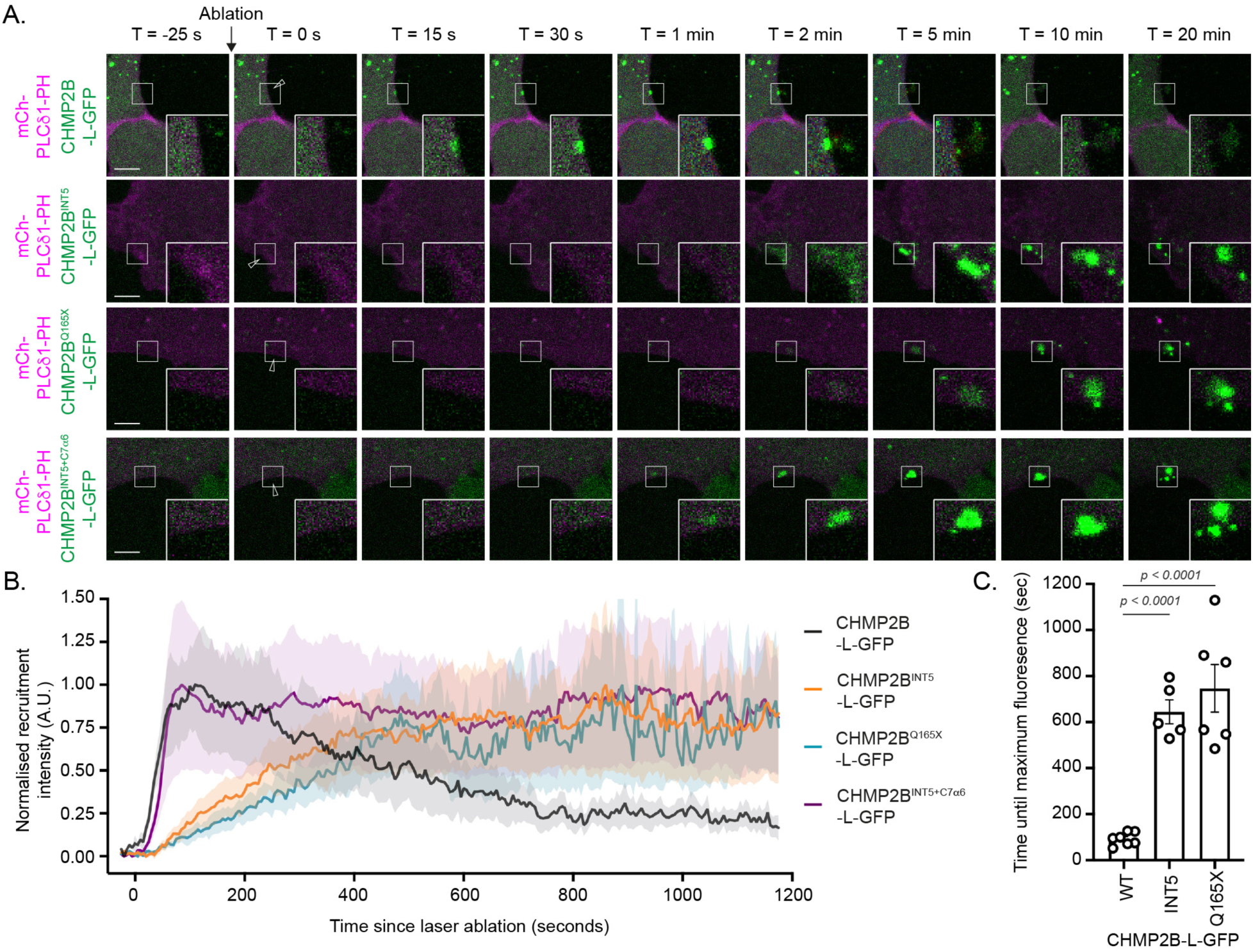
C-truncating variants of CHMP2B arrive late and persist at sites of plasma membrane damage. **A**. Representative image sequences of CAL-51 cells stably expressing CHMP2B-L-GFP, CHMP2B^INT5^-L-GFP, CHMP2B^Q165X^-L-GFP or CHMP2B^INT5+C7NES^-L-GFP and subject to a single multi-photon laser ablation at the mCh-PLC81-PH illuminated plasma membrane. Arrowheads indicate site of laser ablation at T = 0. Full images of cells in this panel are displayed in Figure S3A, and separated channels are displayed in Figure S5. Scale bar is 5 μm. See corresponding movies S1, S3-S5. **B**. Quantification of recruitment dynamics of GFP-fusion proteins in A. Graph displays mean ± S.E.M. from CHMP2B-L-GFP (n = 14 movies, N = 7), CHMP2B^INT5^-L-GFP (n = 11 movies, N = 5), CHMP2B^Q165X^-L-GFP (n = 12 movies, N = 6) or CHMP2B-L-GFP, CHMP2B^INT5+C7NES^-L-GFP (n = 12 movies, N = 6). **C**. Quantification of time to maximum at the site of laser ablation from cells described in B.

The LMB-sensitive nuclear export of CHMP2B-L-GFP and nuclear accumulation of CHMP2A-L-GFP, CHMP2B^INT5^-L-GFP and CHMP2B^Q165X^-L-GFP suggested the presence of as-yet unidentified Nuclear Localisation Sequences (NLS) within these proteins. Using a previously described deletion series of YFP-CHMP2A (Figure S2A and S2B)^48^, we identified a cluster of basic residues in the N-terminus of CHMP2A Helix α2 that allowed nuclear import (Figure S2C - S2F). This arrangement of basic residues is conserved in CHMP2B, and in all other ESCRT-III subunits, (except CHMP1A and CHMP1B (Figure S2G), in which a bi-partite NLS in Helix1 has been described previously^49^). These data highlight a previously unreported difference in subcellular localization between CHMP2A and CHMP2B, explain mechanistically how this is achieved through different NES and NLS activities, and show that FTD-causing mutations in CHMP2B lead to its nuclear retention due to truncation of its NESs.

## C-truncating mutations in CHMP2B display a delayed and persistent recruitment to sites of membrane damage

We next asked whether the altered subcellular distribution of C-truncated FTD mutations of CHMP2B affected its ability to respond to plasma membrane damage. We imaged cells stably expressing CHMP2B-L-GFP live and used a multi-photon laser to ablate a 90 nm^2^ region of the plasma membrane, illuminated with a transfected plasma membrane marker, PLC81^PH^-mCh. Confirming known roles of ESCRT-III in plasma membrane repair, we observed a transient assembly of CHMP2B-L-GFP at the site of damage, reaching a maximum 116 ± 12 seconds after the ablation, before gradually dissipating over the next 20 minutes (Figure 2A, Figure 2B and Figure S3A-S3C, Movie S1). At later stages, we were occasionally able to observe a transient relocation of CHMP2B-L-GFP to an extracellular protrusion and the externalization of plasma membrane at this site (Figure S3B). Interestingly, when the site of ablation was placed at the edge of a filopodia, the dimensionality reduction in these structures allowed us to visualize a ‘wave’ of CHMP2B-L-GFP puncta that formed in the cytosol approximately 20 μm distal to the site of ablation. This wave travelled progressively toward the site of ablation over the next 100 seconds, culminating in the decoration of a patch of membrane which narrowed to a focus at the ablation site (Figure S3C, Movie S2). Whilst prior data has implicated actin remodelling^50^ and lysosomal fusion with the plasma membrane^51^ in plasma membrane repair, we saw no evidence of lysosomes being trafficked to sites of membrane damage, and CHMP2B-L-GFP recruitment prevailed in the presence of actin cytoskeleton disruption (Figure S4). As CHMPs are soluble cytosolic proteins, we infer direct assembly of ESCRT-III at sites of plasma membrane damage. We next examined recruitment of CHMP2B^INT5^-L-GFP and CHMP2B^Q165X^-L-GFP in response to localized plasma membrane damage. Consistent with a limited cytosolic pool of these proteins, both accumulated more slowly at the site of damage, reaching a maximum after 645 ± 52 and 786 ± 107 seconds respectively (Figure 2A-2C, Figure S5, Movie S3 and Movie S4). Whilst recruitment of CHMP2B-L-GFP was transient, recruitment of CHMP2B^INT5^-L-GFP and CHMP2B^Q165X^-L-GFP was persistent, suggesting that turnover of ESCRT-III at sites of membrane damage was impaired (Figure 2A-C). These C-truncating mutations in CHMP2B also remove the MIT-domain Interaction Motif (MIM) present in CHMP2B’s C-terminus that allows engagement of the ESCRT-associated AAA-ATPase, VPS4 (Figure 1E). As VPS4 allows turnover and disassembly of ESCRT-III we wondered whether a failure to bind VPS4 was responsible for the persistence of the C-truncated versions of CHMP2B at sites of membrane damage. Using a stable cell line expressing a chimaera of CHMP2B^INT5^ and the Helix6 NES from CHMP7 (CHMP2B^INT5+C7α6^-L-GFP; Figure S1I) we confirmed that returning this protein to the cytosol restored its timely assembly at sites of plasma membrane damage (Figure 2A, Figure 2B, Figure S5 and Movie S5). Importantly, the NES in CHMP7 Helix6 does not contain a MIM due to charge inversion at a critical arginine residue^45^. Consistent with a role for VPS4 in turning over ESCRT-III, CHMP2B^INT5+C7α6^-L-GFP was not disassembled after recruitment to sites of membrane damage (Figure 2A, Figure 2B and Figure S5). These data separate NES and MIM activities in CHMP2B’s C-terminus, with the NES allowing cytoplasmic access for timely recruitment and the MIM allowing CHMP2B turnover once it has assembled at sites of membrane damage.

## ESCRT-III recruitment is temporally disconnected from membrane sealing

Local Propidium Iodide (PI) influx can be used as a reporter of plasma membrane integrity^22^. We recorded multi-channel live cell imaging data at high time resolution and combined this with multi-photon laser ablation. Laser ablation at the plasma membrane resulted in a burst of PI influx (Figure 3A, Movie S6). As PI entry was transient, rather than progressive, the plasma membrane appears to be almost immediately sealed upon damage. Strikingly, CHMP2B-L-GFP was recruited to sites of damage only after PI influx had abated, suggesting that ESCRT-III recruitment is disconnected from the act of membrane sealing (Figure 3B). We next examined PI influx in cells expressing CHMP2B^INT5^-L-GFP or CHMP2B^Q165X^-L-GFP. Consistent with ESCRT-III recruitment occurring after membrane sealing, the kinetics of PI influx in these cells was indistinguishable from WT cells, despite CHMP2B^INT5^-L-GFP or CHMP2B^Q165X^-L-GFP localizing to sites of damage substantially later than CHMP2B-L-GFP (Figure 3B). These data indicate that membrane sealing, and ESCRT-III recruitment are temporally disconnected and suggest that ESCRT-III-independent factors are responsible for the initial seal of damaged membranes.

**Figure 3.**
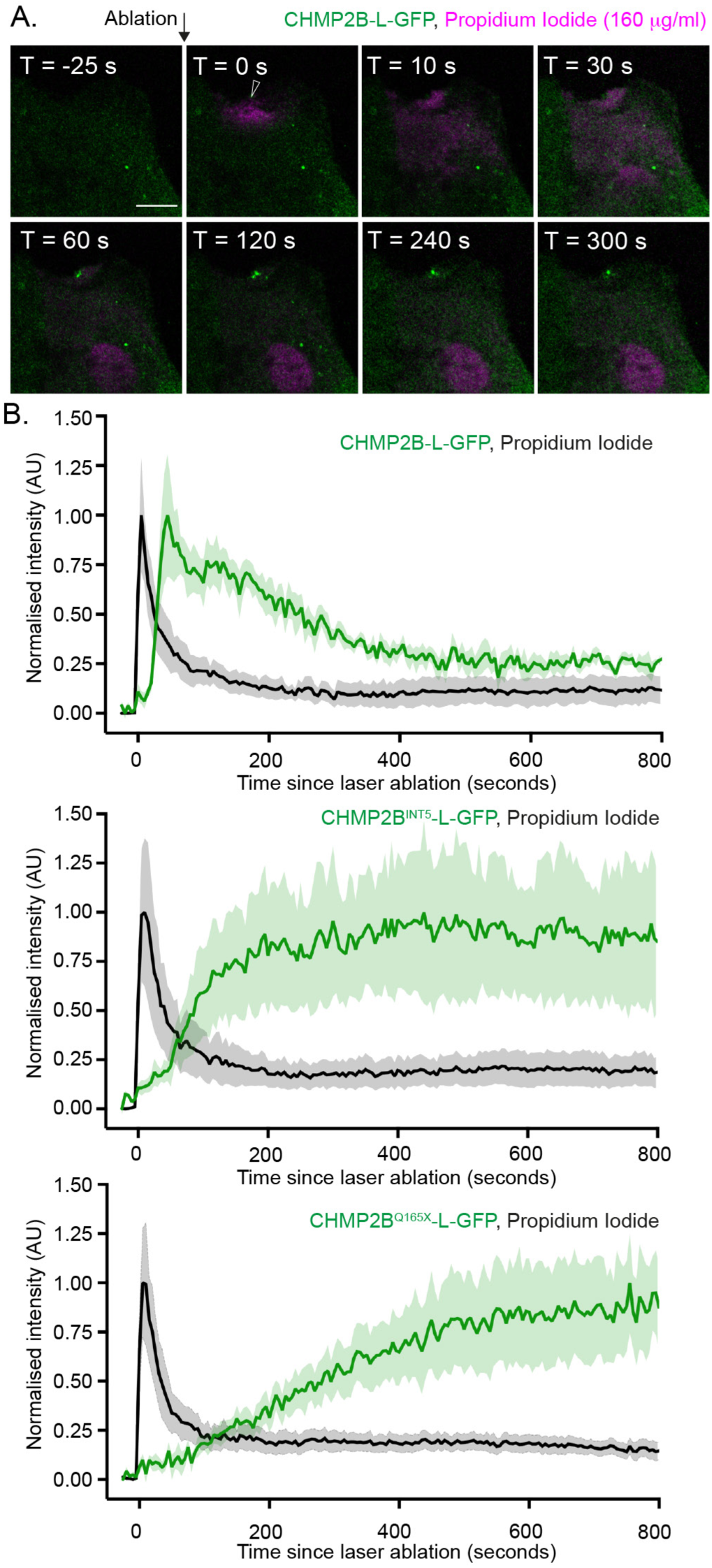
CHMP2B-L-GFP arrives at sites of membrane damage after membranes have been sealed. **A**. Representative image sequences of CAL-51 cells stably expressing CHMP2B-L-GFP and subject to a single multi-photon laser ablation in the presence of external Propidium Iodide (PI, 160 μg/mL, magenta). Arrowhead indicates site of laser ablation at T = 0. Scale bar is 10 μm. See corresponding movie S6. **B**. Quantification of PI influx and recruitment dynamics of CHMP2B-L-GFP (n = 15 movies, N = 5), CHMP2B^INT5^-L-GFP (n = 8 movies, N = 4), CHMP2B^Q165X^-L-GFP (n = 12 movies, N = 7). Graph displays mean ± S.E.M.

## Annexin proteins are recruited immediately to sites of plasma membrane damage in advance of ESCRT-III

As CHMP2B was recruited to sites of plasma membrane damage after membranes had been sealed, we next looked upstream at potential recruitment factors for ESCRT-III. Human cells express a total of 11 Annexin proteins (Figure S6A) that bind both calcium and phospholipid membranes. ANXA7 and ANXA11 are unique in that they share an extended N-terminal low-complexity domain (LCD) that binds the ESCRT-associated factor, Apoptosis-linked gene-2 (ALG-2)^52,53^ and participates in liquid-liquid phase separation^54^. To investigate coordination between ESCRT-III and these factors, we generated stable cell lines expressing both CHMP2B-L-GFP and mCherry (mCh) tagged versions of these proteins. Building on the discovery that ALG-2 and ANXA7 were recruited immediately to sites of plasma membrane damage,^55^, we used multi-photon laser ablation of the plasma membrane, to confirm that ALG-2 was enriched progressively at the point of damage, peaking just before CHMP2B-L-GFP and forming projections that protruded out of the cell (Figure 4A, Movie S7). Like ANXA7-mCh (Movie S8), we found that ANXA11-mCh, was recruited immediately (ANXA11-mCh; 5.9 ± 0.5 seconds) to sites of damage, first decorating a broad area of the plasma membrane and rapidly becoming focused to a point at the site of ablation that projected out of the cell (ANXA11-mCh; 9.5 ± 2.9 seconds after ablation) (Figure 4B and 4C, Movie S9). CHMP2B-L-GFP was recruited some 40 seconds later (45.3 ± 3.1 seconds after ablation), initially covering a broad area of the plasma membrane surrounding the Annexin focus (Figure 4B and 4C) with its intensity peaking 82 ± 8.3 seconds after ablation. Importantly, ANXA11-mCh recruitment was temporally coordinated with the block in local PI influx, suggesting that Annexin recruitment plays a role in the sealing of damaged membranes (Figure 4D).

**Figure 4.**
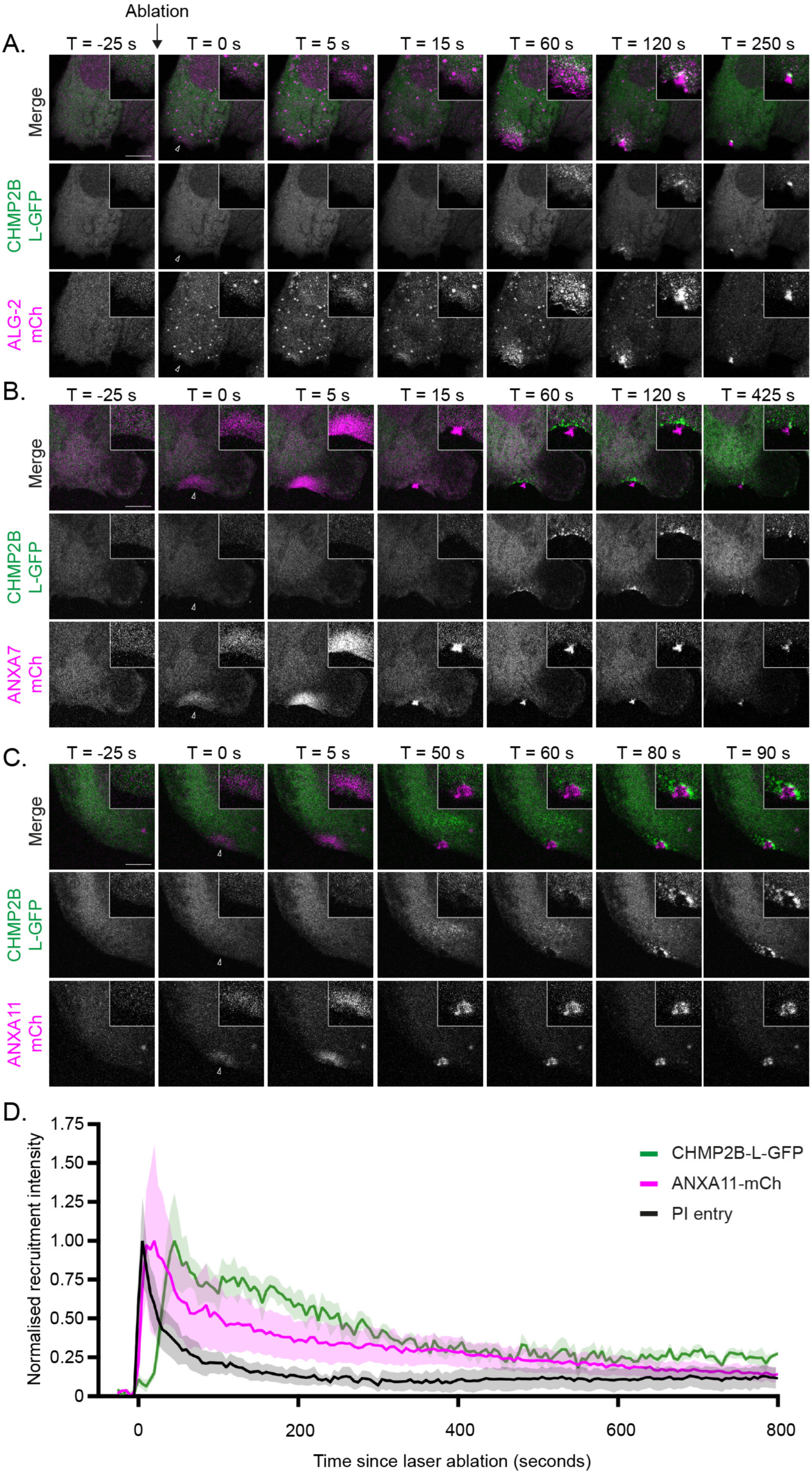
ALG-2, ANXA7 and ANXA11 are recruited to sites of membrane damage in advance of CHMP2B. **A-C.** Representative image sequences of CAL-51 cells stably expressing CHMP2B-L-GFP and either ALG-2-mCh (n = 5, N = 2), ANXA7-mCh (n = 8, N = 3) or ANXA11-mCh (n = 11, N = 3) and subject to a single multi-photon laser ablation at the plasma membrane. Number of movies in each case indicated. Arrowheads indicate site of laser ablation at T = 0. Scale bar is 10 μm. See corresponding movies S7-S9. **D**. Overlay of PI influx dynamics and CHMP2B-L-GFP and ANXA11-mCh recruitment dynamics downstream of a single multi-photon laser ablation. The average ANXA11-mCh (n = 5, N = 3) was overlaid on the CHMP2B-L-GFP and PI traces from Figure 3B. Data displayed as mean ± S.E.M.

## ALS-associated mutations in ANXA11 impair its recruitment to sites of membrane damage

Pathogenic mutations in ANXA11 (including G38R, R235Q, R346C and R456H), are associated with ALS^40,56^, a neurodegenerative disease found on a continuum with FTD. The P93S mutation was recently found to be a pathogenic variant in corticobasal syndrome^57^. To test whether mutations in ANXA11 impacted its ability to respond to membrane damage, we examined recruitment of WT, several pathogenic mutants and a LCD-deletion of ANXA11-mCh to laser-ablated lesions of the plasma membrane. ANXA11-mCh, ANXA11^G38R^-mCh, ANXA11^P93S^-mCh and ANXA11^R346C^-mCh were recruited immediately to the site of damage and concentrated rapidly at a focus that protruded from the cell (Figure 5A and 5B, Figure S6B, Movie S10 – Movie S13). In contrast to this, ANXA11^8LCD^-mCh, ANXA11^R235Q^-mCh and ANXA11^R456H^-mCh did not assemble at sites of laser ablation in the plasma membrane (Figure 5A and Figure 5B, Figure S6B, Movie S14 - S16). In all cases, CHMP2B-L-GFP was recruited to the site of damage approximately 40 seconds after ablation (Figure 5C), first surrounding the Annexin cluster and then narrowing to a focus at the neck of this protrusion. These data suggest that the LCD is essential for ANXA11 clustering at sites of membrane damage, and that ALS-causing mutations in ANXA11 can impair this process.

**Figure 5.**
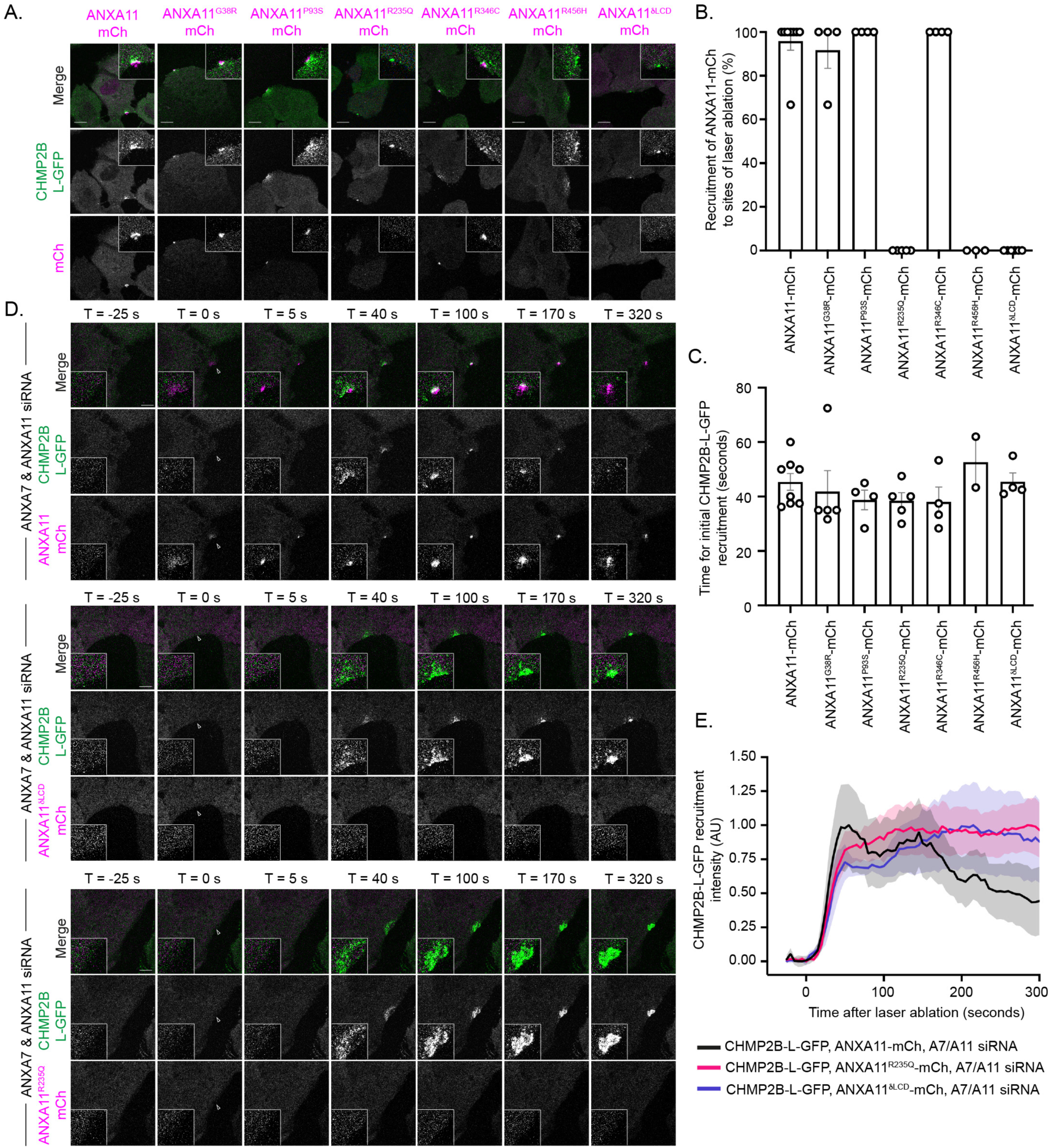
Pathogenic ALS-associated mutations in ANXA11 are not recruited to sites of membrane damage. **A**. Representative images of CAL-51 cells stably expressing CHMP2B-L-GFP and either ANXA11-mCh, ANXA11^G38R^-mCh, ANXA11^P93S^-mCh, ANXA11^R235Q^-mCh, ANXA11^R456H^-mCh or ANXA11^8LCD^-mCh and subject to a single multi-photon laser ablation. Recruitment of CHMP2B-L-GFP (enlarged in boxed region) was used as a positive indicator of the membrane damage response. Image sequences from these stills displayed in Figure S6B. Scale bar is 10 μm. See corresponding movies S10-S16. **B**. Frequency of wildtype and mutant ANXA11-mCh recruitment to sites of laser-ablation from A. Graph displays mean ± S.E.M. CHMP2B-L-GFP:ANXA11-mCh (n = 23, N = 8); CHMP2B-L-GFP:ANXA11^G38R^-mCh (n = **9**, N = 4); CHMP2B-L-GFP:ANXA11^P93S^-mCh (n = 11, N = 4); CHMP2B-L-GFP:ANXA11^R235Q^-mCh, n = 12 cells, N = **5**; CHMP2B-L-GFP:ANXA11^R346C^-mCh, n = 12 cells, N = 4; CHMP2B-L-GFP:ANXA11^R456H^-mCh, n = 11 cells, N = 3; CHMP2B-L-GFP:ANXA11^8LCD^-mCh, n = 16 cells, N = 6. **C**. Quantification of CHMP2B-L-GFP recruitment time in cells expressing the indicated ANXA11-mCh proteins. Graph displays mean ± S.E.M. from the indicated number of independent experiments. CHMP2B-L-GFP:ANXA11-mCh (n = 21, N = 8); CHMP2B-L-GFP:ANXA11^G38R^-mCh (n = 10, N = 5); CHMP2B-L-GFP:ANXA11^P93S^-mCh (n = 12, N = 4); CHMP2B-L-GFP:ANXA11^R235Q^-mCh (n = 9, N = 5); CHMP2B-L-GFP:ANXA11^R346C^-mCh (n = 11, N = 4); CHMP2B-L-GFP:ANXA11^R456H^-mCh (n = 8, N = 2); CHMP2B-L-GFP:ANXA11^8LCD^-mCh (n = 13, N = 4). No significant differences in CHMP2B-L-GFP recruitment time, calculated by one-way ANOVA, were observed across these cell lines. **D**. Representative image sequences from CAL-51 cells treated with siRNA-targeting ANXA7 and ANXA11, stably expressing CHMP2B-L-GFP and either siRNA resistant ANXA11-mCh, ANXA11^R235Q^-mCh or ANXA11^8LCD^-mCh, and subject to a single multi-photon laser ablation at the plasma membrane, indicated by the arrowhead. Scale bar is 5 μm. **E**. Quantification of recruitment and turnover of CHMP2B-L-GFP in the cell lines described in D. Mean ± S.E.M. displayed from ANXA7/ANXA11 siRNA treated CHMP2B-L-GFP:ANXA11-mCh (n = 10, N = 4); CHMP2B-L-GFP:ANXA11^R235Q^ -mCh, (n = 11, N = 4); CHMP2B-L-GFP:ANXA11^8LCD^-mCh (n = 14, N = 4). See corresponding movies S17-S19.

## ALS-causing mutations in ANXA11 impair ESCRT-III turnover at sites of plasma membrane damage

Given the similarities in recruitment to sites of membrane damage of the LCD-containing Annexins ANXA7 and ANXA11, we next used siRNA to deplete endogenous ANXA7 and ANXA11 from cells stably expressing CHMP2B-L-GFP and re-expressed siRNA-resistant versions of ANXA11-mCh to investigate requirements for ANXA11 in the response to membrane damage. In ANXA7 and ANXA11-depleted cells re-expressing siRNA-resistant ANXA11-mCh, CHMP2B-L-GFP was recruited on schedule to the site of damage and was brought to a focus at the cell-proximal side of the ANXA11-mCh-positive protrusion (Figure 5D, Figure 5E and Movie S17). In contrast, siRNA-resistant ANXA11^8LCD^-mCh or ANXA11^R235Q^-mCh, were no longer assembled at the site of membrane damage (Figure 5D). Strikingly, while CHMP2B-L-GFP was still recruited on schedule in these cells (Figure 5D, Figure 5E and Figure S7A), it did not focus to a point (Figure S7B) and instead persisted at the site of damage (Figure 5D, Figure 5E, Movie S18 and Movie S19). The persistence of CHMP2B-L-GFP at sites of plasma membrane damage in the presence of ANXA11^8LCD^-L-mCh and ANXA11^R235Q^-L-mCh mimicked that seen with CHMP2B^INT5^-L-GFP or CHMP2B^Q165X^-L-GFP, suggesting that in the presence of these pathogenic versions of ANXA11, ESCRT-III activity at the site of membrane damage is stalled. These data suggest that impaired ESCRT-III turnover at sites of membrane damage may be a common feature of these forms of FTD and ALS.

## Pore-forming toxins allow investigation of a population-level plasma membrane damage response

To extend our findings with a more physiological insult, we generated recombinant versions of the *Clostridium perfringens* cholesterol dependent cytolysin, Perfringolysin-O (PfO). We exposed cells to 1 nM PfO in the presence of external PI. Consistent with established oligomerization and pore-formation times, we observed local PI entry into cells after 10-15 minutes. As with laser-ablation, after PfO treatment, PI entry across the plasma membrane was transient, again appearing as a burst (Figure 6A, Figure 6B and Movie S20). These data suggest that once assembled and permeable, the PfO pore is rapidly sealed, presumably through an acute mechanism to restrict cytolysis. We examined CHMP2B-L-GFP and again found recruitment to the plasma membrane at sites of PI influx, but only after the incoming burst of PI had dispersed (Figure 6A and Figure 6B, Movie S20). These data suggest that across different insults, cells mount an acute response to seal damaged membranes and that in both cases, ESCRT-III components are recruited after this seal has been formed.

**Figure 6.**
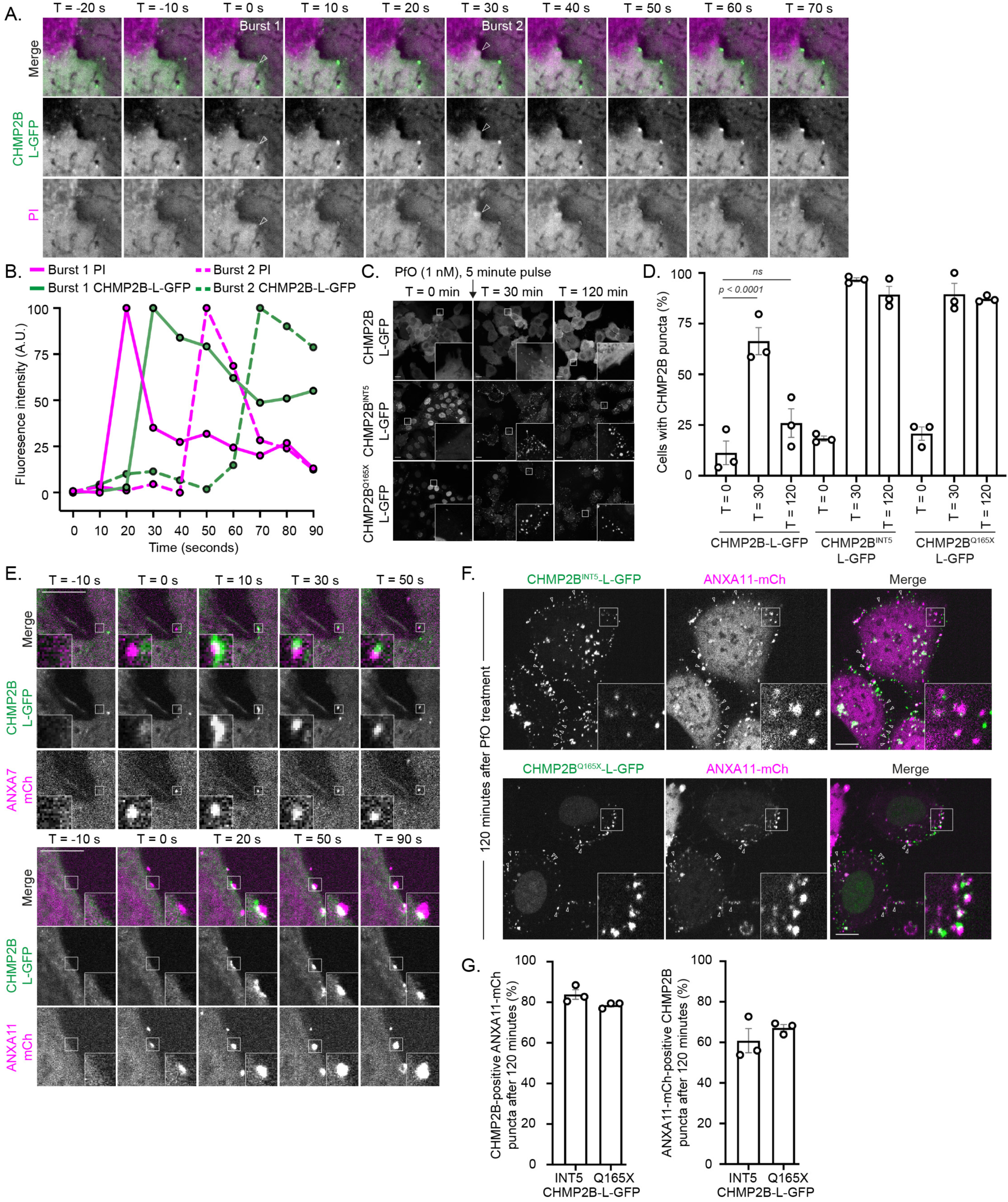
Sequential recruitment of ANXA7/ANXA11 and CHMP2B-L-GFP to sites of PfO-mediated membrane damage. **A**. Representative image sequence of CAL-51 cells stably expressing CHMP2B-L-GFP and treated with 1 nM PfO in the presence of 160 μg/mL PI. Bursts of cytoplasmic PI entry marked with arrowheads; note recruitment of CHMP2B-L-GFP in frames following PI entry, and that PI entry is transient, rather than progressive. See corresponding movie S20. **B**. Quantification of GFP and PI fluorescent intensities in the cytoplasmic region of Burst-1 and Burst-2 from A. **C**, **D**. CAL-51 cells stably expressing CHMP2B-L-GFP, CHMP2B^INT5^-L-GFP or CHMP2B^Q165X^-L-GFP were treated with 1 nM PfO for 5 minutes and chased into fresh media for the indicated times. Cells displaying GFP-puncta at the plasma membrane at the indicated times were scored manually; graph displays mean ± S.E.M. from between 353 and 617 cells per condition analysed across N = 3 independent experiments. Significance calculated by one-way ANOVA, with Tukey correction. **E**. CAL-51 cells stably expressing CHMP2B-L-GFP and either ANXA7-mCh or ANXA11-mCh were treated with 1 nM PfO for 5 minutes and then chased into fresh media and imaged live. Sequential recruitment of ANXA7-mCh/ANXA11-mCh and CHMP2B-L-GFP to cell-distal and cell-proximal regions of the protrusion was observed in 90/94 ANXA7 recruitment events from N = 5 independently imaged cells (ANXA7-mCh, CHMP2B-L-GFP) and from 107/109 events from N = 4 independently imaged cells (ANXA11-mCh, CHMP2B-L-GFP). Scale bar is 10 μm. Timestamp set relative to Annexin recruitment. See corresponding movies S21 and S22. **F**, **G**. CAL-51 cells stably expressing CHMP2B^INT5^-L-GFP or CHMP2B^Q165X^-L-GFP and ANXA11-mCh were treated with 1 nM PfO for 5 minutes, chased into fresh media and imaged live after 120 minutes. Graph (G) displays mean ± S.E.M. of colocalization or juxtaposition of CHMP2B-L-GFP and ANXA11-mCh signals from 2663 puncta present in 100 cells (CHMP2B^INT5^-L-GFP) or 2107 puncta present in 84 cells (CHMP2B^Q165X^-L-GFP) displaying GFP-puncta at the plasma membrane and imaged across N = 3 independent experiments. Representative images (F) with additional examples of colocalisation displayed by arrowheads.

In continual presence of external PfO, multiple sites of membrane damage led to substantial PI influx across cells in the population, resulting eventually in saturation of PI in the nucleus. We used this phenotype to examine mutations in PfO that were predicted to prevent its cytolytic activity whilst retaining a wildtype threshold for cholesterol binding (PfO^Y181A/F318A^)^58^ or through impairment of monomer-monomer interactions required for pre-pore to pore transition (PfO^K336E^)^59^. We verified that these mutations prevented both PfO-induced PI influx (Figure S8A-C) and the recruitment of CHMP2B-L-GFP at the plasma membrane (Figure S8D), linking CHMP2B recruitment to the pore-forming activity of the toxin. We next examined the persistence of CHMP2B-L-GFP, CHMP2B^INT5^-L-GFP or CHMP2B^Q165X^-L-GFP at damage sites induced by a 5-minute pulse of PfO. As with laser ablation, whilst CHMP2B-L-GFP assembled transiently, CHMP2B^INT5^-L-GFP and CHMP2B^Q165X^-L-GFP persisted at the sites of PfO-mediated membrane damage (Figure 6C, Figure 6D, Figure S8E, Figure S8F, Movie S21 – S23). Using doubly stable cell lines, we also confirmed that ANXA7-mCh/ANXA11-mCh and CHMP2B-L-GFP were recruited sequentially to sites of PfO-mediated membrane damage (Figure 6E). As with our laser ablated cells, ANXA7-mCh and ANXA11-mCh accumulated on the cell-distal side of membrane protrusions, with CHMP2B-L-GFP recruited later and being focused to a point at the cell-proximal side of these protrusions (Figure 6E, Movie S24 and Movie S25). Outwardly budded structures have been observed previously by scanning electron microscopy at the site of laser ablation^22^ and our data suggest similar structures are formed in the response to pore forming toxin-mediated membrane damage, with a common spatiotemporal segregation of membrane repair factors within these protrusions. Given the impaired turnover of CHMP2B^INT5^-L-GFP and CHMP2B^Q165X^-L-GFP at sites of plasma membrane damage (Figure 6C and Figure 6D), we next examined whether these puncta retained Annexins. In cells treated with PfO for 5 minutes and imaged two hours later, we found that the ANXA11-mCh puncta were retained at the CHMP2B^INT5^-L-GFP and CHMP2B^Q165X^-L-GFP accumulations at sites of plasma membrane damage (Figure 6F and 6G). We confirmed that endogenous ANXA11 was similarly retained at the persistent CHMP2B^INT5^-L-GFP and CHMP2B^Q165X^-L-GFP puncta at the plasma membrane 2 hours after PfO-mediated plasma membrane damage, whereas it was largely cleared from the plasma membrane in cells expressing CHMP2B-L-GFP (Figure S9). These data link the impaired turnover of the FTD-associated C-truncated CHMP2B mutants at sites of plasma membrane damage to the retention of ANXA11 at these locations.

## Damaged membranes are shed from cells

ESCRT-III acts to release enveloped retroviruses such as HIV-1 from the plasma membrane by severing the membrane neck connecting budding virions to the plasma membrane. Prior work studying MLKL and GSDMD pores in the context of necroptosis and pyroptosis respectively has observed ESCRT-III localisation to the plasma membrane protrusions^15,60^ with the inference that these protrusions are to be shed. Additionally, calcium dependent shedding of Streptolysin-O-containing material has been previously observed by centrifugal capture^61^, although a mechanistic basis for this shedding is lacking. Given the observed localisation of CHMP2B-L-GFP to the base of ANXA11-mCh-positive protrusions formed after PfO-mediated membrane damage, and known roles for ESCRT-III in reverse topology membrane scission^27^, we wondered if we could detect shedding of these protrusions from cells. During timelapse imaging of PfO-treated cells stably expressing ANXA7-mCh and CHMP2B-L-GFP, or ANXA11-mCh and CHMP2B-L-GFP, we could detect the abrupt disappearance of the mCh-Annexin protrusion in the frames after CHMP2B-L-GFP recruitment, suggesting that Annexin-positive protrusion is shed from cells (Figure 7A, Movie 26 and Movie 27). We observed sequential recruitment of Annexins and CHMP2B-L-GFP, with the ANXA7-mCh or ANXA11-mCh-positive protrusion lost within 2 minutes of CHMP2B-L-GFP recruitment (Figure 7B). To confirm the release of PfO and Annexin-positive membranes biochemically, we treated cells with a sub-lytic pulse of PfO for 5 minutes, washed the monolayer extensively, chased the cells into fresh media (Figure 7C) and collected material released from cells by filtration and ultracentrifugation. We found that PfO could be recovered from the media (Figure 7D and Figure 7E) and found that recovery of pelletable ANXA7, ANXA11, ALIX and ALG-2 was elevated in the media of cells treated with PfO (Figure 7D and Figure 7E). Importantly, both release of PfO, and enhanced release of ANXA7, ANXA11, ALG-2 and the Annexin, ALG-2 and ESCRT-III-binding protein, ALIX, was not observed in cells treated with PfO^Y181A/F318A^, which does not form pores (Fig S8A-C). This links ESCRT-III recruitment (Figure S8D) and shedding of these membranes directly to the membrane damaging activity of this toxin (Figure 7D and Figure 7E).

**Figure 7:**
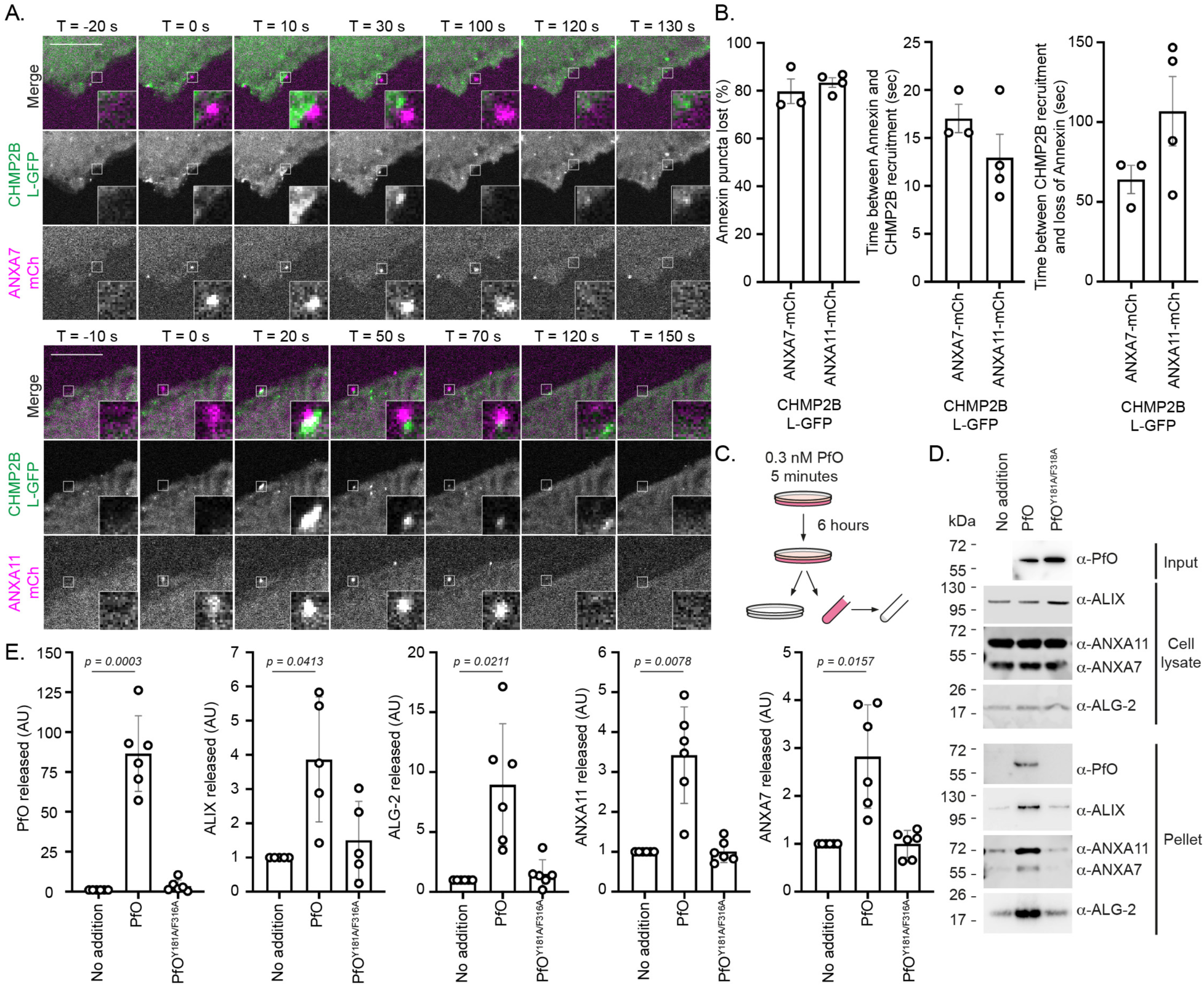
Annexin-containing membranes are shed after PfO-mediated membrane damage. **A.** Representative images of CAL-51 cells stably expressing CHMP2B-L-GFP and either ANXA7-mCh or ANXA11-mCh that had been treated with a 5-minute pulse of 1 nM PfO, chased into fresh media and imaged live. T = 0 was set at the time of ANXA7-mCh or ANXA11-mCh puncta formation. See corresponding movies S23 and S24. **B**. Quantification of recruitment dynamics and release of ANXA7-mCh or ANXA11-mCh puncta in the first 5 minutes of imaging in A. Loss of ANXA positive puncta and time between ANXA7-mCh or ANXA11-mCh and CHMP2B-L-GFP recruitment (ANXA7-mCh puncta; mean ± S.E.M. from 61 events across 3 independent experiments; ANXA11-mCh; mean ± S.E.M. from 44 events across 4 independent experiments) and the time between CHMP2B-L-GFP recruitment and the loss of the ANXA7-mCh or ANXA11-mCh puncta (ANXA7-mCh puncta loss; mean ± S.E.M. from 47 events across 3 independent experiments; ANXA11-mCh; mean ± S.E.M. from 37 events across 4 independent experiments) was scored. No significant differences between ANXA7-mCh and ANXA11-mCh expressing cell lines were observed, as calculated by unpaired two-tailed T-test. **C**. Schematic for PfO-dependent shedding assay. Briefly, cells were treated for with 0.3 nM PfO or PfO^Y181A/F318A^ for 10 minutes and chased into fresh media for 6 hours. Media was collected and subject to ultracentrifugation to capture released material. **D**. Resolved cell lysates and pelleted media fractions were examined by western blotting with antisera raised against PfO, ALIX, ANXA7, ANXA11 or ALG-2. **E**. Quantification of band intensity from D. All values were normalized to “no addition” control signal. Graphs display mean ± S.D. from N = 5 (ALIX) or N = 6 (all others) independent experiments with significance calculated by one-way ANOVA with Dunnett’s correction. Scale bar is 10 μm.

## In the absence of ESCRT-activity, plasma membrane seals are leaky

As the response to membrane damage involves immediate Annexin recruitment, a rapid seal and a subsequent shedding of Annexin-positive material, we wondered if the ANXA7/ANXA11 assembly represented a temporary ‘seal’ to preserve membrane integrity, followed by an ESCRT-III-dependent shedding of this membrane to ‘heal’ the phospholipid bilayer. We investigated the robustness of this seal by treating cells with a 5-minute pulse of PfO and probing plasma membrane integrity by adding PI at sequential times after removal of PfO (Figure 8A). At 30 minutes after PfO addition, CHMP2B-L-GFP, CHMP2B^INT5^-L-GFP or CHMP2B^Q165X^-L-GFP cells were all equally susceptible to PI challenge (Figure 8B and Figure 8C). Allowing a chase of 120-minutes before challenging with external PI revealed that cells expressing CHMP2B-L-GFP were no longer permeable to PI. However, at this time point, cells expressing CHMP2B^INT5^-L-GFP or CHMP2B^Q165X^-L-GFP were still able to allow PI entry (Figure 8B and Figure 8C), indicating that the plasma membrane integrity of these cells was still impacted by PfO. Cells expressing CHMP2B^INT5^-L-GFP or CHMP2B^Q165X^-L-GFP were also able to let more PI in over this period of chase, leading to significant nuclear accumulation of the dye (Figure 8D). We next wondered if the persistent loss of membrane integrity in cells expressing CHMP2B^INT5^-L-GFP or CHMP2B^Q165X^-L-GFP led to cytotoxicity. We exposed cells to differing concentrations of PfO and followed cell death by live Incucyte microscopy. We found that cells expressing CHMP2B^INT5^-L-GFP or CHMP2B^Q165X^-L-GFP were more sensitive to PfO than cells expressing CHMP2B-L-GFP (Figure 8E). These data suggest that whilst cells mount an acute response to membrane damage by mobilising Annexin proteins, this initial seal is not robust and requires ESCRT-III activity to normalise and heal these membranes through a process of shedding. Moreover, these data show that cells bearing pathogenic mutations of CHMP2B are less able to maintain membrane integrity and more susceptible to cell death upon plasma membrane damage.

**Figure 8:**
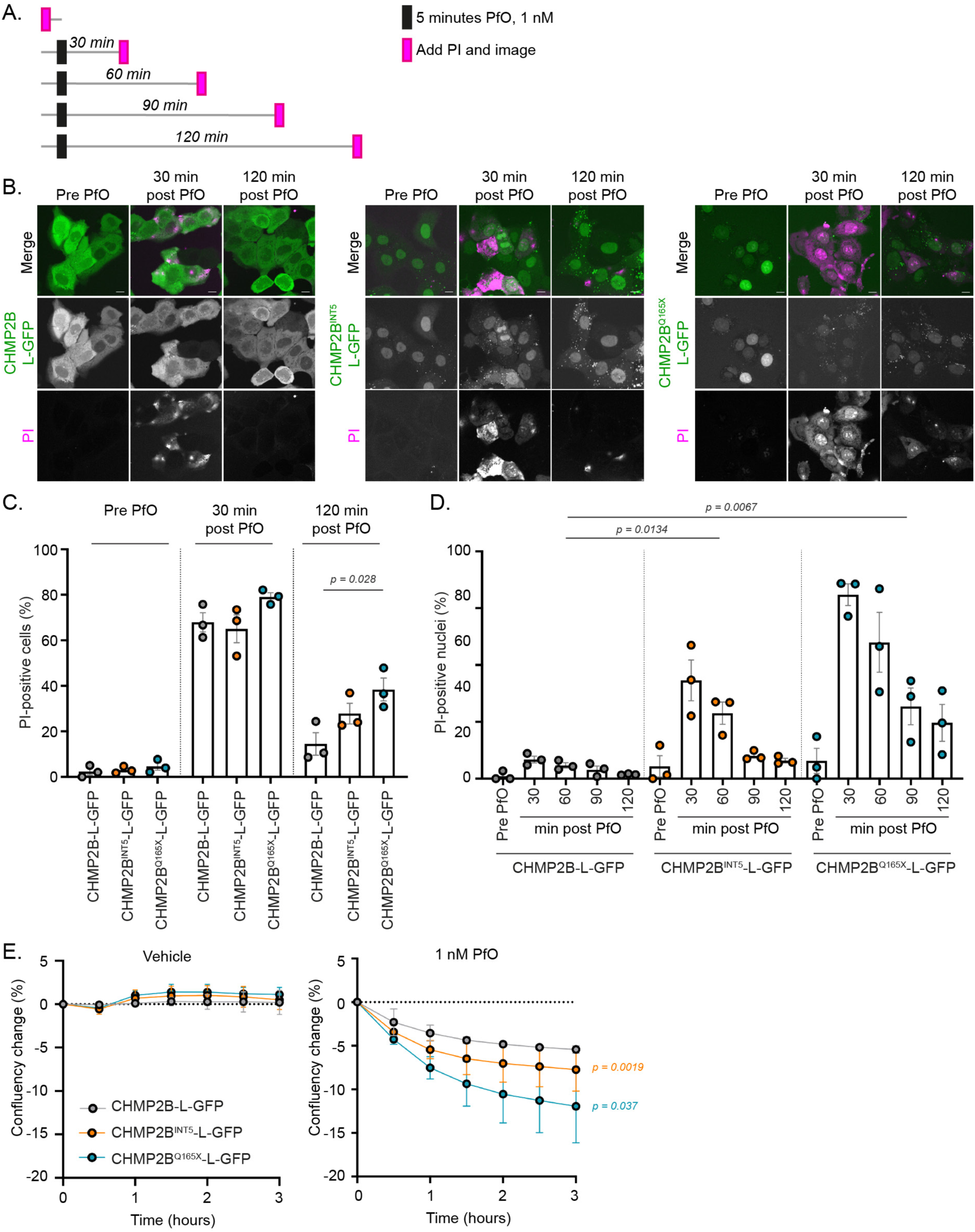
Cells bearing FTD-causing mutations in CHMP2B are more sensitive to membrane damaging agents. **A**. Schematic of membrane damage protocol to examine time-dependent membrane repair. **B**-**D**. Representative images (B) and quantification (C, D) of CAL-51 cells stably expressing CHMP2B-L-GFP, CHMP2B^INT5^-L-GFP, CHMP2B^Q165X^-L-GFP, treated with 1 nM PfO for 5 minutes, chased into fresh media and challenged with PI at the indicated timepoints. Scale bar in (B) is 10 μm. The number of PI positive cells (C) and the number of cells with PI-positive nuclei (indicative of saturation) (D) was quantified from 2784 cells (CHMP2B-L-GFP) 2268 cells (CHMP2B^INT5^-L-GFP) and 1951 cells (CHMP2B^Q165X^-L-GFP) from N = 3 independent experiments. Mean ± S.E.M. displayed with statistical significance determined by unpaired T-test (C), two-way ANOVA across all timepoints and cell lines (D). **E**. CAL-51 cells stably expressing CHMP2B-L-GFP, CHMP2B^INT5^-L-GFP, CHMP2B^Q165X^-L-GFP were treated with the indicated concentration of PfO and imaged live using Incucyte microscopy. Cytotoxicity led to a reduced confluency of the monolayer and was calculated from N = 3 independent experiments, with three technical repeats per experiment and normalised to T = 0. Mean ± S.E.M. displayed with significance calculated by one-way ANOVA across all timepoints.

## Discussion

Here, we have used live cell imaging to illuminate how Annexin and ESCRT-III proteins work together in the cellular response to plasma membrane damage. We have uncovered nuanced stages in the repair process; an acute seal that limits loss of cellular material, and a subsequent shedding process that restores the membrane. Several machineries have been proposed to orchestrate the repair of damaged membranes, although it is unclear whether these represent independent mechanisms of membrane repair, or whether each machinery represents a single facet of a multi-stage process. Here, using independent forms of plasma membrane damage, we show that arrival of the calcium-binding ANXA7 or ANXA11 proteins at sites of damage occurs concurrently with the initial sealing of the plasma membrane. One possibility is that the accumulation of Annexin proteins at these sites, in a LCD-dependent manner, acts as an acute plug to restrict cytolysis and facilitate subsequent ESCRT-III-dependent repair. Interestingly, this could have parallels with the condensation of stress granules that act to seal damaged endolysosomes^20^, the phase separation of LEM2’s LCD that acts to seal holes in the nuclear envelope^62^, the ESCRT-III-independent diffusion barriers that maintain nucleocytoplasmic compartmentalization prior to ESCRT-III activity during *S. pombe* spindle pole body extrusion^63^ and the finding that biomolecular condensates can contribute to the membrane bending necessary for intra-endosomal vesicle formation^64^. We found that ESCRT-III proteins were only recruited to sites of damage after Annexin recruitment, and after permeability to PI had been restricted, indicating that ESCRT-III recruitment occurs after membranes have been sealed. Consistent with this, in a recent cryo-electron tomography study of perforin-mediated cytolysis cellular membranes appear intact at the point of ESCRT-III recruitment^65^. Overexpressed CHMP2B has been shown previously to assemble at plasma membrane protrusions in cells^66^ and to the base of membrane necks pulled from giant unilamellar vesicles, where it acted as a diffusion barrier to restrict lipid flow^67^. We speculate that this activity might isolate damaged membrane lipids and the Annexin plug for subsequent ESCRT-III-dependent shedding. Prior data has observed membrane protrusions at sites of laser ablation^22^ and shown that pore forming toxins could be collected from the media of treated cells^61,68^. Whilst ESCRT-III proteins have been reported to localise to the plasma membrane of cells damaged by MLKL or GasderminD pores^15,60^, the process of shedding these membranes at this site has only been inferred. Here, we show that ANXA7 and ANXA11 proteins are involved in the coordination of these processes by condensing rapidly into protrusions at sites of membrane damage that were removed from the cell after ESCRT-III recruitment.

These data highlight topological parallels in the shedding of Annexin-containing membranes with the release of enveloped retroviruses, during which the act of ESCRT-III-dependent budding leaves a scarless plasma membrane. Additionally, the dynamic relocalisation of CHMP2B from a broad initial recruitment to a focus at the neck of an ANXA7-or ANXA11-positive protrusion parallels the relocalisation of ESCRT-III during cytokinetic abscission^69,70^, suggesting that the behaviour of ESCRT-III is conserved between these processes. Whilst Annexin and ESCRT-III proteins clearly work together, recruitment of CHMP2B was independent of ANXA11, suggesting that independent localisation cues act to mobilise these proteins. However, mimicking the phenotype of C-truncated versions of CHMP2B, in the absence of ANXA11 recruitment, CHMP2B accumulated under the site of damage, rather than condensing transiently to the neck of an ANXA11-positive protrusion. These data suggest that condensation of ANXA11 may help shape the membrane into a neck for subsequent ESCRT-III dependent severing. Our data suggests a segregation of sequential ‘sealing’ and ‘healing’ activities between Annexin and ESCRT-III proteins during membrane repair.

These defects in membrane repair were uncovered through the analysis of FTD- and ALS-causing mutations in the ESCRT-III subunit CHMP2B and ANXA11. Whilst the C-truncated pathogenic versions of CHMP2B have been well characterized in terms of an endolysosomal phenotype, reflective of impaired ability to recruit VPS4 through C-terminal MIMs^71,72^, we note here relative differences in their subcellular localization. Similar differences in the subcellular localisation of CHMP2A and CHMP2B were observed, with CHMP2A predominantly localised to the nucleus and CHMP2B to the cytoplasm. These data suggest that functional segregation across the Vps2 paralogs may have been enabled by gene duplication within ESCRT-III. We identify functional NESs that drive CHMP2B’s cytoplasmic localization and show that they are lost in the C-truncated FTD-associated versions of this protein. As such, as well as plasma membrane repair, we hypothesise that the kinetics of a range of cytosolic ESCRT-III activities such as endolysosomal repair and multivesicular body biogenesis are likely compromised by these pathogenic mutants of CHMP2B. Increasing evidence presents FTD/ALS as a spectrum of disease, inclusive of a wide range of genetic risk factors. Thus far, there is no consensus as to an underlying mechanism supporting this relationship. Here, we have demonstrated that two FTD/ALS associated proteins act sequentially in the same process of plasma membrane repair. Moreover, pathogenic mutations in both CHMP2B and ANXA11 converge on a common phenotype of impaired ESCRT-III turnover at sites of membrane repair, suggesting membrane damage as a potential disease mechanism in FTD/ALS.

## Acknowledgements

We thank Dr Kurt Anderson and Dr Matt Renshaw (Crick Advanced Light Microscopy) for calibration assistance for the laser ablation assays. We thank Prof. Mark Wallace (Dept. Chemistry, King’s) for advice relating to pore forming toxins.

## Materials and Methods

### Cell Culture

STR-profiled, mycoplasma-free vials of GP2-293 (CVCL_WI48), Phoenix-AMPHO (CVCL_H716) and CAL-51 (CVCL_1110) cells were obtained from the Crick Cell Services Science Technology Platform. Cells were cultured at 37 °C and at 5% CO_2_ in Dulbecco’s Modified Eagle Medium (DMEM) containing 10% FBS, penicillin (100 U/mL) and streptomycin (0.1 mg/mL). For live cell imaging, DMEM was exchanged for FluoroBrite DMEM (Gibco) with identical supplementation and 2 mM L-glutamine.

### Plasmids

cDNAs encoding coding sequences for CHMP2B, CHMP2B^INT^^5^ or CHMP2B^Q165X^ were synthesised with flanking 5’ *EcoRI* and 3’ *NotI* sites and cloned into the *EcoRI* and *NotI* sites of a modified version of the retroviral expression vector pCMS28^73^ bearing a EGFP-based Localisation and Affinity Purification tag^44^ C-terminal to the *NotI* site to create the vector series pCMS28 CHMP2B-L-GFP, pCMS28 CHMP2B^INT^^5^-L-GFP and pCMS28 CHMP2B^Q165X^-L-GFP.

CHMP2B coding sequences were transferred to pCR3.1 *EcoRI*-*XhoI*-*NotI*-HA by subcloning. Codon optimised and RNAi-resistant coding sequences for ANXA7 and ANXA11 were synthesised with 5’ *EcoRI* and 3’ *NotI* sites and cloned into the *EcoRI* and *NotI* sites of a modified version of pMSCVneo bearing a sequence encoding mCherry C-terminal to the *NotI* site. A plasmid encoding Perfringolysin-O (PfO) was a kind gift from Prof. Alejandro Heuck (Univ. Massachusetts). PfO sequences were amplified with flanking *EcoRI* and *NotI* sites and cloned into the *EcoRI* and *NotI* sites of pET28a (Novagen). Mutations to all plasmids were created by standard Polymerase Chain Reaction-based techniques. Plasmids were verified by Sanger sequencing (GeneWIZ). A plasmid encoding MP-Act-mRuby3 was a kind gift from Prof Tobias Meyer (Stanford) via Addgene (155221). A plasmid encoding LAMP1-mCh was a kind gift from Dr Max Gutierrez (Crick).

### Recombinant Protein Production

*Escherichia coli* BL21(DE3) cells expressing pET28a-PfO plasmids were grown at 37 °C to O.D. 0.6 in LB medium, induced with 0.25 mM IPTG for 3 hours, harvested by centrifugation and resuspended in TNG500 lysis buffer (50 mM Tris-HCl pH 7.5, 500 mM NaCl, 10% w/v Glycerol) supplemented with cOmplete Protease Inhibitors (Roche), 10 mM imidazole, 5 mM β-Mercaptoethanol, 0.2% Triton X-100, 1 mM PMSF, 10 μg/mL DNaseI, and 1 mg/mL lysozyme. Resuspended cells were lysed by sonication and clarified by centrifugation at 40,000 x g for 60 minutes. Supernatants were incubated with Ni-NTA resin for 2 hours at 4 °C. The resin was washed sequentially with TNG500 lysis buffer, then with TNG500 lysis buffer containing 0.01% Triton X-100 and four steps of this buffer containing increasing imidazole concentrations (20, 30, 40, 50 mM imidazole) with corresponding reductions in NaCl concentrations to preserve osmolarity. Proteins were eluted in elution buffer (50 mM Tris-HCl pH 7.5, 100 mM NaCl, 400 mM imidazole, 5 mM β-Mercaptoethanol, 10% w/v Glycerol, 0.01% Triton X-100), dialysed into TNG150 (50 mM Tris-HCl pH 7.5, 500 mM NaCl, 10% w/v Glycerol) and assessed by SDS-PAGE and western blotting to assess purity. Bio-Rad Protein Assay was used to assess concentration.

### Antibodies and Chemical Reagents

Anti-GFP (clone 7.1/13.1) was from Roche; in-house anti-CHMP2B (3335) was developed by BioGenes as previously described^74^; anti-GAPDH (clone 6C5, MAB374) was from Millipore; anti-ALIX (12-422-1-AP), anti-ANXA7 (10154-2-AP), anti-ANXA11 (10479-2-AP) and anti-ALG-2 (12303-1-AP) were from Proteintech; anti-PfO (ab225685) was from Abcam; anti-p42/p44 MAPK (9102) was from Cell Signalling Technology; anti-HIS (652502, clone J099B12) was from Biolegend. Alexa conjugated secondary antibodies were from Invitrogen, HRP-conjugated secondary antibodies (7074 and 7076) were from Cell Signalling Technology. IRDye 800 CW (925-32210) and IRDye 680 RD (925-68071) were from LI-COR Biosciences. Propidium Iodide was from Merck. Leptomycin B was from Cayman Chemicals. Cytochalasin D was from Merck.

### Transient transfection of cDNA

CAL-51 cells were transfected using Lipofectamine-3000 (Life Technologies) according to the manufacturer’s instructions. GP2-293 cells were transfected using linear 25-kDa polyethylenimine (PEI, Polysciences, Inc.), as described previously^73^.

### siRNA Depletion

All siRNA-based depletions were performed at 20 nM final concentration using Lipofectamine RNAiMAX (Life Technologies) according to the manufacturer’s instructions. siRNA were obtained from Horizon Discovery. Control siRNA was D-001810-01. ANXA7 was depleted with J-010760-08 (CCGAGAAAUUGUCAGAUGU), ANXA11 was depleted with J-011212-05 (GAAGAUCUGUGGUGGCAAU).

### Retrovirus generation

GP2-293 or Pheonix-AMPHO cells were transfected with retroviral expression vectors and pVSVG (Takara). Viral supernatants were harvested 48 hours later, purified by low-speed centrifugation and 0.2 μm syringe filtration and added to target cells in the presence of 0.8 μg/mL polybrene. Selective antibiotics (Puromycin, 1 μg/mL; G418 0.8 mg/mL) were applied after 48 hours.

### Fixation and Immunofluorescence

Cells were plated on 13 mm #1.5 coverslips and fixed in 4 % PFA for 20 minutes at room temperature, then washed thrice in PBS. For staining, fixed cells were permeabilized using 0.2 % Triton-X100, washed thrice in PBS, quenched with 0.2 M glycine and blocked in 1 % BSA for 30 minutes. Cells were incubated with primary antibodies (1:200) in 1 % BSA for 2 hours at room temperature, washed thrice with PBS before incubation with secondary antibodies in 1 % BSA for 1 hour at room temperature. Coverslips were washed and mounted onto microscope slides with fluorescent mounting medium (Dako) and imaged using an Andor Dragonfly 200 spinning disc confocal paired with a Zyla 5.5 sCMOS camera and using a Nikon Eclipse Ti2 with Plan Apo 60x/1.4NA or 100x/1.45NA objectives.

### Live Cell Imaging and Laser Ablation Microscopy

Cells expressing the indicated proteins were plated in 8-well μslides (Ibidi) approximately 24 hours prior to imaging. Live imaging was performed using the Andor Dragonfly 200 spinning disc confocal, as described above, with an environmental chamber (Okolabs) to maintain temperature at 37 °C and 5% CO_2_. For laser ablation assays cells were imaged with a Zeiss LSM 880 imaging system in a Tokai Hit stage top incubator at 37 °C, 5 % CO_2_. Images were acquired with a 60X oil immersion objective every 5 seconds. After the first 5 frames, cells were ablated at a manually defined area of 0.09 μm^2^ (1 x 1 pixel) with a tuneable Chameleon OPO laser (Coherent) at 800 nm with 15-20 % laser power. At the beginning of each imaging experiment, a test ablation was performed to ensure optimal membrane targeting. The collimator position was then adjusted to alter the z-plane as required. In experiments investigating propidium iodide (PI) influx, cells were preincubated for 10 minutes with 160 μg/mL PI in FluoroBrite DMEM. For quantification of laser ablation experiments, regions of interest were drawn around the maximal zone of CHMP2B-L-GFP recruitment and projected across time points. Mean intensity values were measured in these ROIs for each frame. Corresponding background values were measured similarly and used for frame-wise background correction by subtraction. For PI measurements, a region of interest was drawn around the local area of influx at T = 0 and integrated density was applied and measured across all frames. Movies with significant drift, retraction of cells after membrane damage, or death were discarded from the analysis.

### Incucyte Microscopy

CAL-51 cells stably expressing CHMP2B-L-GFP, CHMP2B^INT5^-L-GFP or CHMP2B^Q165X^-L-GFP were seeded in Phenoplate glass bottomed 96-well plates (Perkin Elmer). PfO (1 nM) was diluted in complete imaging media and imaged using a Sartorius Incucyte SX5 at 37 °C and 5% CO2, capturing 9 images per well every 30 minutes. Sartorius Incucyte Analysis software was used to calculate the average confluency in each well normalised to T = 0 and acquired in triplicate for each independent experiment.

### Collection of released material by ultracentrifugation

CAL-51 cells stably expressing the indicated proteins were grown in two 15 cm dishes per condition in complete DMEM. When confluent, cells were washed thrice with PBS and then into DMEM containing 1% FBS. PfO was added at a final concentration of 0.3 nM in warmed DMEM containing 1% FBS for 10 minutes. PfO-containing media was removed, the monolayer was washed thrice with PBS, taking care to remove all residual PBS, and cells were incubated with 18 mL of DMEM containing 1% FBS. Cells were left at 37 °C for 6 hours to allow PfO-containing membranes to be released. The media was collected and subject to low-speed centrifugation (300 x g, 5 minutes) to pellet any insoluble material, and was filtered through a 5 μm PluriStrainer. 36 mL of media per condition was subject to ultracentrifugation at 100,000 x g for 6 hours at 4 °C in a SW-32-Ti swinging bucket rotor (Beckman). The media was decanted and 500 μL PBS was added to the pellet and left to resuspend overnight at 4 °C. The next morning, the pellet was resuspended and re-centrifuged at 100,000 x g in low-bind 1.5 mL ultracentrifuge tubes using a TLA55 fixed angle rotor (Beckman). The pellet was resuspended in 50 μL of PBS and denatured through the addition of 50 μL 2 x LDS sample buffer containing 100 mM DTT.

### Statistical analysis

Two-tailed Student’s T-tests, or ordinary one-way ANOVA with the indicated post-hoc tests, were used to assess significance between test samples and controls or significance in response over time, as stated in figure legends. N-numbers given as the number of independent experiments; n-numbers given as the number of cells or images analysed.

---

**Figure S1.**
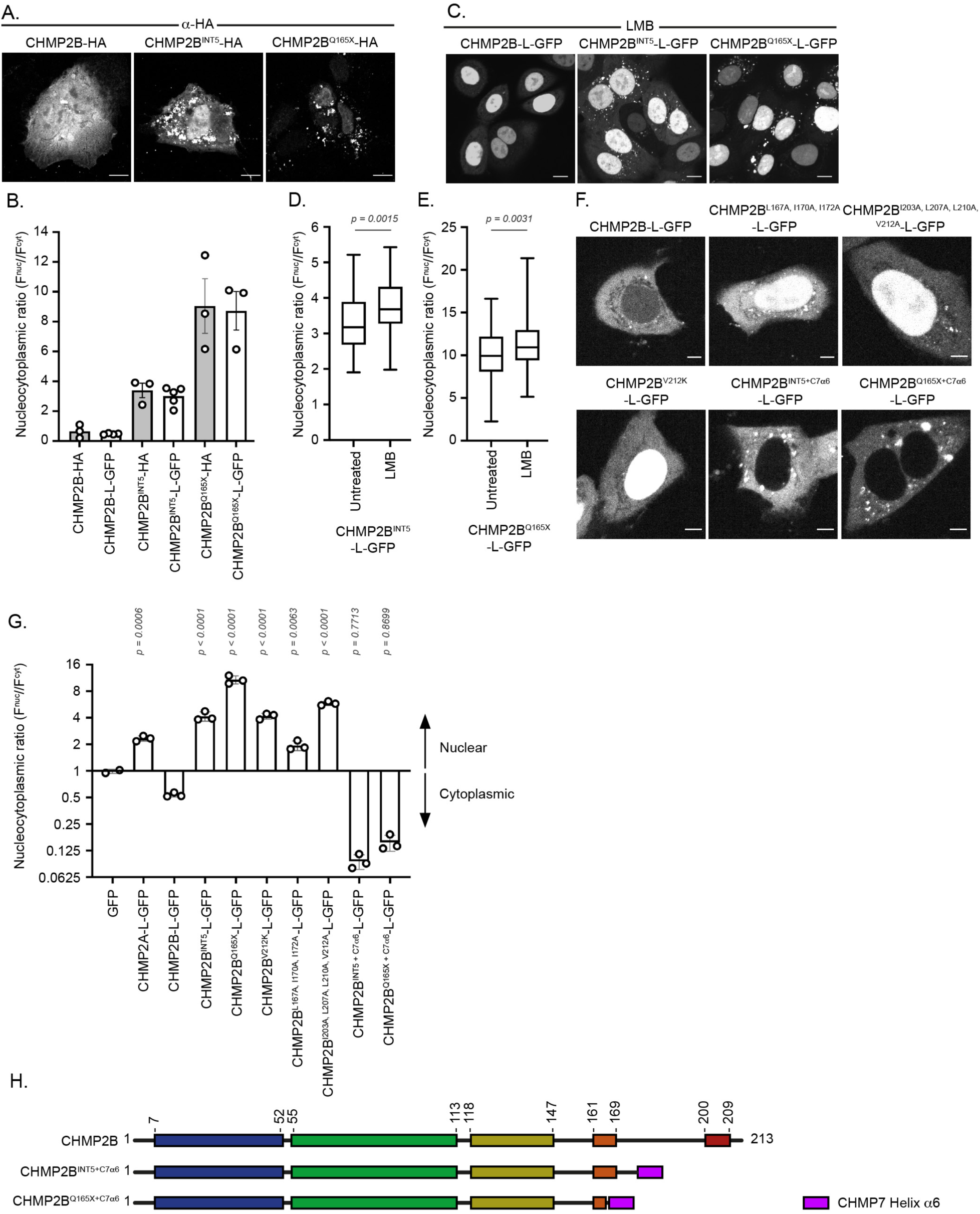
CHMP2B encodes C-terminal nuclear export sequences that are truncated by pathogenic FTD-causing mutations. **A**. Representative images of CAL-51 cells transiently transfected with HA-tagged CHMP2B and illuminated with antisera raised against HA. Scale bars represent 10 µm. **B.** Quantification of nucleocytoplasmic ratio of HA-tag fluorescence from cells in A or compared to the CAL-51 cell lines stably expressing equivalent CHMP2B-L-GFP fusion proteins (data as shown in Fig.1B). CHMP2B-HA (n = 25), CHMP2B^INT5^-HA (n = 30), CHMP2B^Q165X^-HA (n = 26). CHMP2B-L-GFP (n = 57), CHMP2B^INT5^-L-GFP (n = 31), CHMP2B^Q165X^-L-GFP (n = 105). For HA-tagged versions, data was combined from N = 3 independent experiments. For GFP-tagged versions, data was combined from N = 3-5 independent experiments as indicated. Graphs show mean ± S.E.M. with significance between HA vs. L-GFP equivalent cells calculated by two-tailed T-test. **C**. Representative images of CAL-51 cells stably expressing CHMP2B-L-GFP, CHMP2B^INT5^-L-GFP or CHMP2B-L-GFP^Q165X^ and treated with LMB (10 ng/mL, 4 hours). Scale bars represent 10 µm. **D, E**. Quantification of nucleocytoplasmic partitioning of cells expressing CHMP2B^INT5^-L-GFP and treated with LMB (n = 36 (untreated) or 73 (+LMB)) or CHMP2B-L-GFP^Q165X^ (n = 72 (untreated) or 85 (+LMB)), related to Figure 1D. Data displayed as a box and whisker plot with median, 25^th^ and 75^th^ percentile displayed. Significance calculated by 2-tailed T-test. . **F**. Representative live images of CAL-51 cells transiently transfected with CHMP2B-L-GFP, (n = 41), CHMP2B^L167A/I170A/I172A^-L-GFP, (n = 46), CHMP2B^I203A/L207A/L210A/V212A^-L-GFP (n = 56), CHMP2B^V212K^-L-GFP, (n = 67), CHMP2B^INT5+C7α6^-L-GFP, (n = 42), CHMP2B^Q165X+C7α6^-L-GFP (n = 55) from N = 3 independent experiments. Scale bars represent 5 μm. **G**. Quantification of nucleocytoplasmic ratio of GFP fluorescence of cells described in (F). Graph displays mean ± S.D. from N = 3 independent experiments (N = 2 for GFP) with significance calculated by one-way ANOVA. **H**. Schematic depicting chimaeric sequences of CHMP2B^INT5^ and CHMP2B^Q165X^ fused with sequences from CHMP7 Helix6 containing a type-1 NES. Helix boundaries depicted by alternating colours and placed according to Alphafold-2 predictions of CHMP2B (Q9UQN3). Addition of CHMP7’s NES sequence indicated by magenta box.

**Figure S2.**
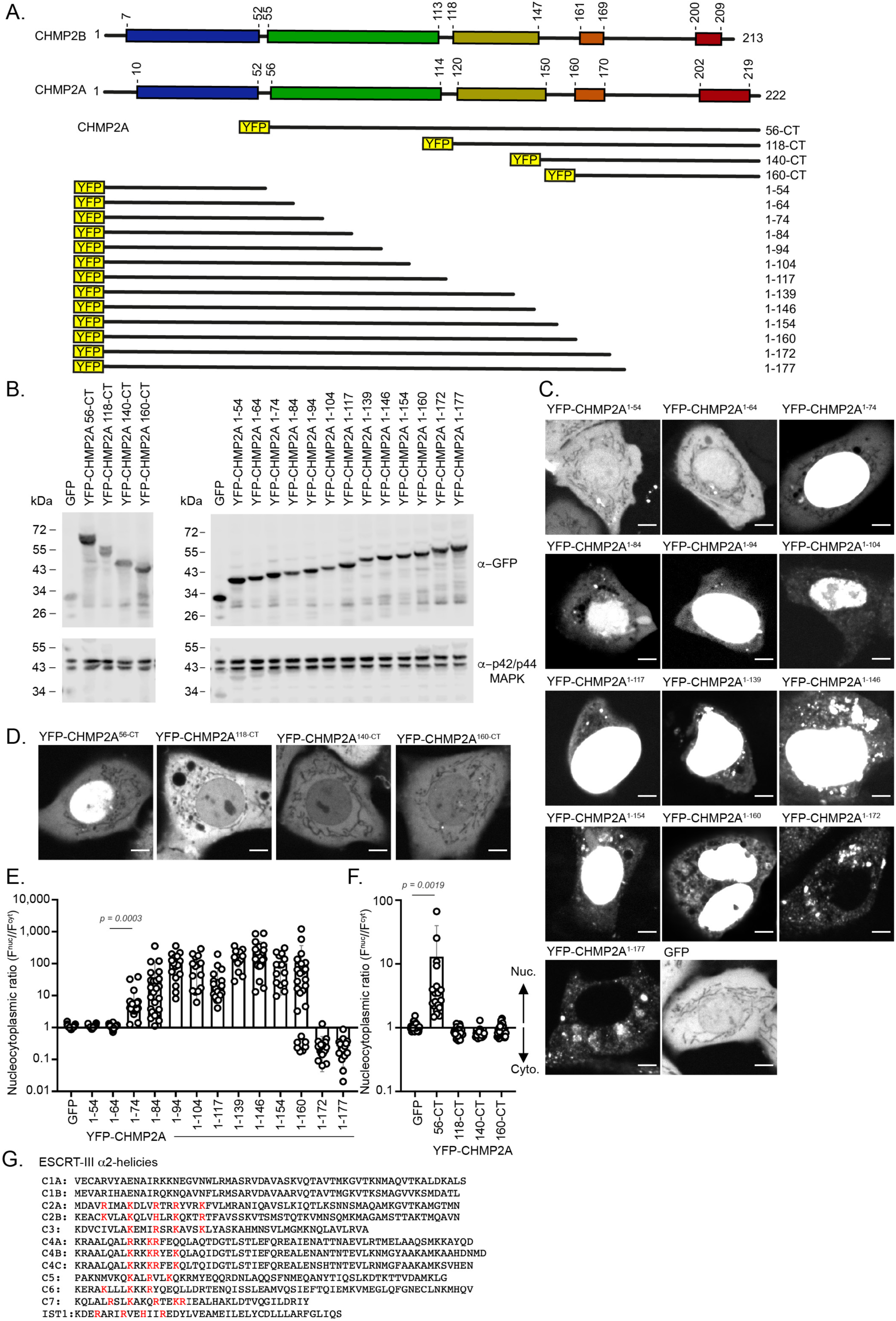
Conserved nuclear localisation sequences in the α2 helix of ESCRT-III proteins. **A.** Schematic representation of YFP-CHMP2A deletions (described in^48^). **B**. Resolved lysates from 293T cells transfected with vectors encoding GFP, or the constructs from A were examined by western blotting with antisera raised against GFP or p42/p44 MAPK. **C**, **D**. Representative images of CAL-51 cells transiently transfected with GFP or the constructs depicted in (A) and imaged live. Scale bars represent 5 μm. **E, F**. Quantification of nucleocytoplasmic ratio of YFP fluorescence from CAL-51 cells transfected with GFP or the constructs described in A. Graphs show mean ± S.E.M. from n = 15-51 cells per condition across N = 1 independent experiment with significance calculated by two tailed T-test. Error bars below zero cannot be displayed on a logarithmic y-axis. **G**. Sequence alignment of α2 helices in mammalian ESCRT-III proteins. A cluster of basic residues at the N-terminus of these helices highlighted in red.

**Figure S3.**
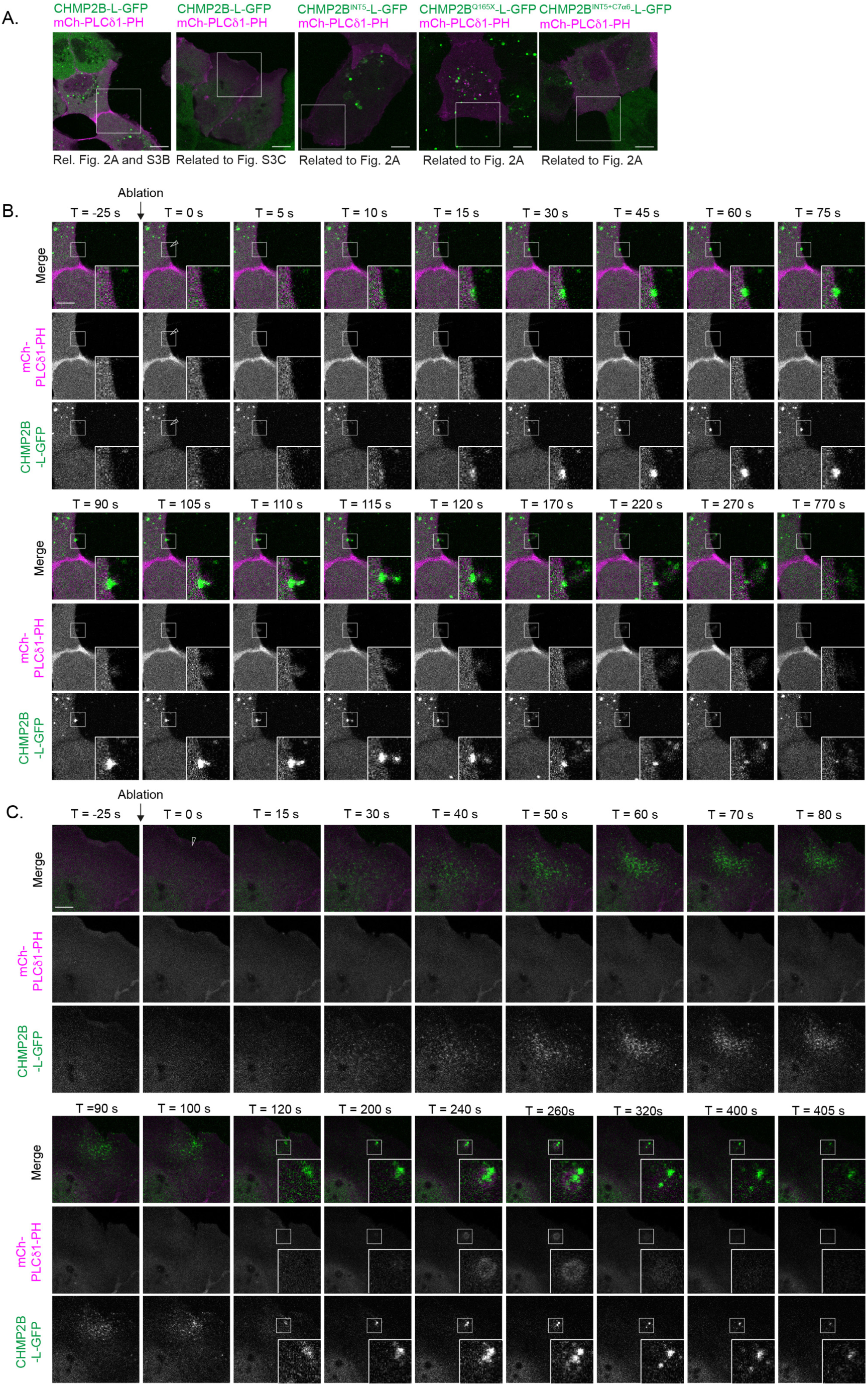
Characterisation of CHMP2B-L-GFP recruitment dynamics at sites of laser ablation. **A**. Whole cell images of cropped image sequences presented in Figure 2A, Figure S3B, Figure S3C and Figure S5. Images taken from frames prior to laser ablation. Scale bar is 10 μm. **B, C.** Representative images of CAL-51 cells stably expressing CHMP2B-L-GFP, transiently transfected with a vector encoding PLCδ1-PH-mCh to illuminate the plasma membrane and subject to a single multi-photon laser ablation at the plasma membrane. In (B), additional frames from the image sequence in Figure 2A are presented. Here, CHMP2B-L-GFP recruitment appears as a single punctum, which gradually disappears. As depicted, we were occasionally able to visualise events that seemed to be the budding of CHMP2B-L-GFP from this site. In C, the site of laser ablation was placed at the edge of a lamellipodium, enabling the visualisation of CHMP2B-L-GFP recruitment in a stellate wave-form pattern focusing on the site of damage, with CHMP2B-L-GFP being focused at the site of damage. In B & C, Scale bar is 5 μm. See corresponding movies S1 and S2.

**Figure S4.**
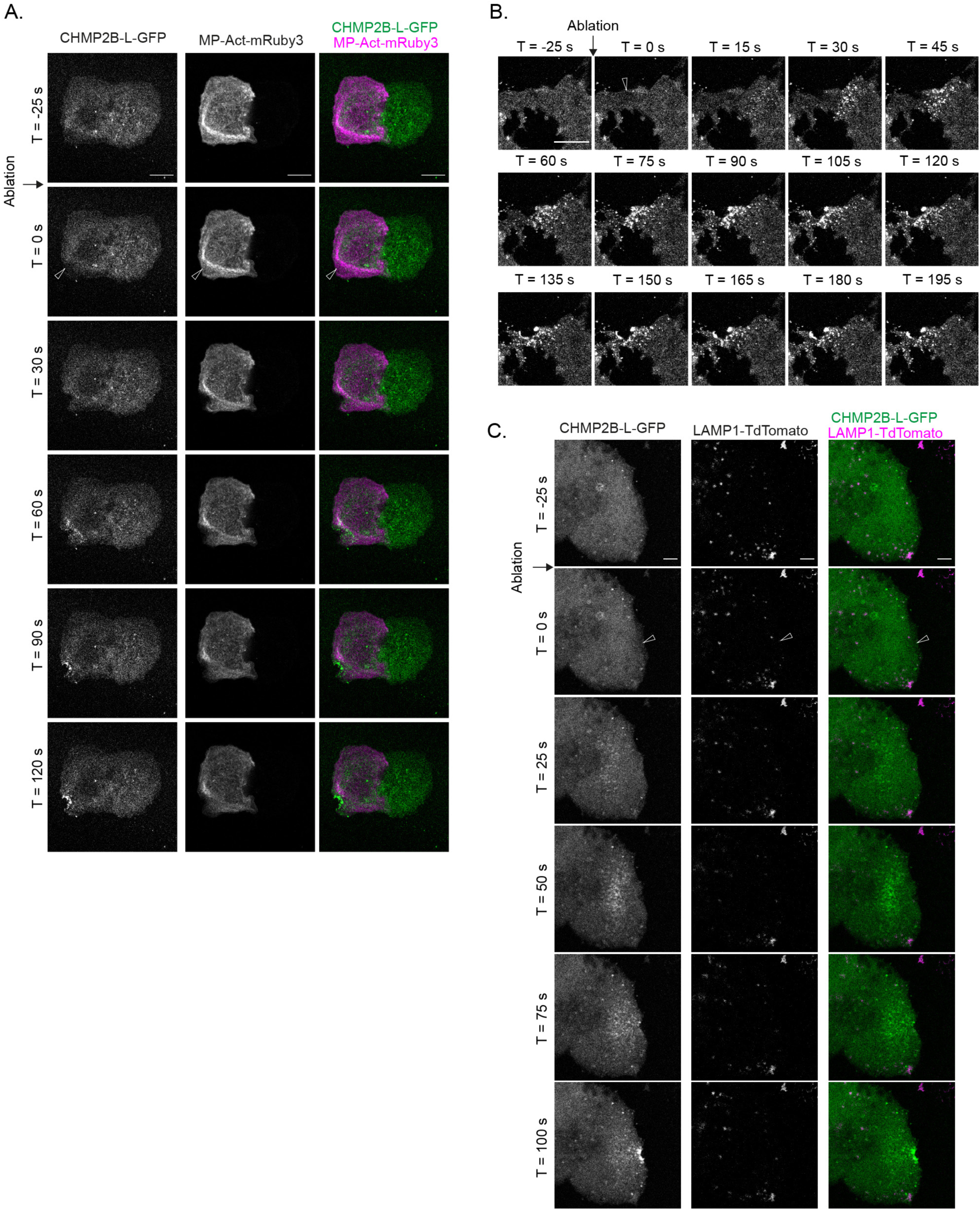
Recruitment of CHMP2B-L-GFP to sites of membrane damage occurs independently of the actin cytoskeleton and does not require lysosomes. **A**. Representative image sequence of CAL-51 cells stably expressing CHMP2B-L-GFP and transiently transfected with MPAct-mRuby3 to illuminate membrane proximal F-actin and subject to a single multi-photon laser ablation at the plasma membrane. Scale bars represent 10 μm. White arrow indicates area of ablation. Images representative of n = 5 cases where clear CHMP2B-L-GFP recruitment was observed, but with no MPAct-mRuby3 relocalisation. See corresponding movie S25. **B**. Representative image sequence from CAL-51 cells stably expressing CHMP2B-L-GFP, pretreated with cytochalasin D (2 μg/mL, 30 minutes prior) and subject to a single multi-photon laser ablation at the plasma membrane. Scale bars represent 10 μm. Arrowhead indicates area of ablation. Images representative of n = 7 cases where clear CHMP2B-L-GFP recruitment was observed. See corresponding movie S26. **C**. Representative image sequence from CAL-51 cells stably expressing CHMP2B-L-GFP, transiently transfected with a plasmid encoding LAMP1-tdTomato and subject to a single multi-photon laser ablation at the plasma membrane. Images representative of n = 7 movies where CHMP2B-L-GFP recruitment, but not LAMP1-mCh recruitment, was observed. Scale bars represent 5 μm. White arrow indicates area of ablation. See corresponding movie S27.

**Figure S5.**
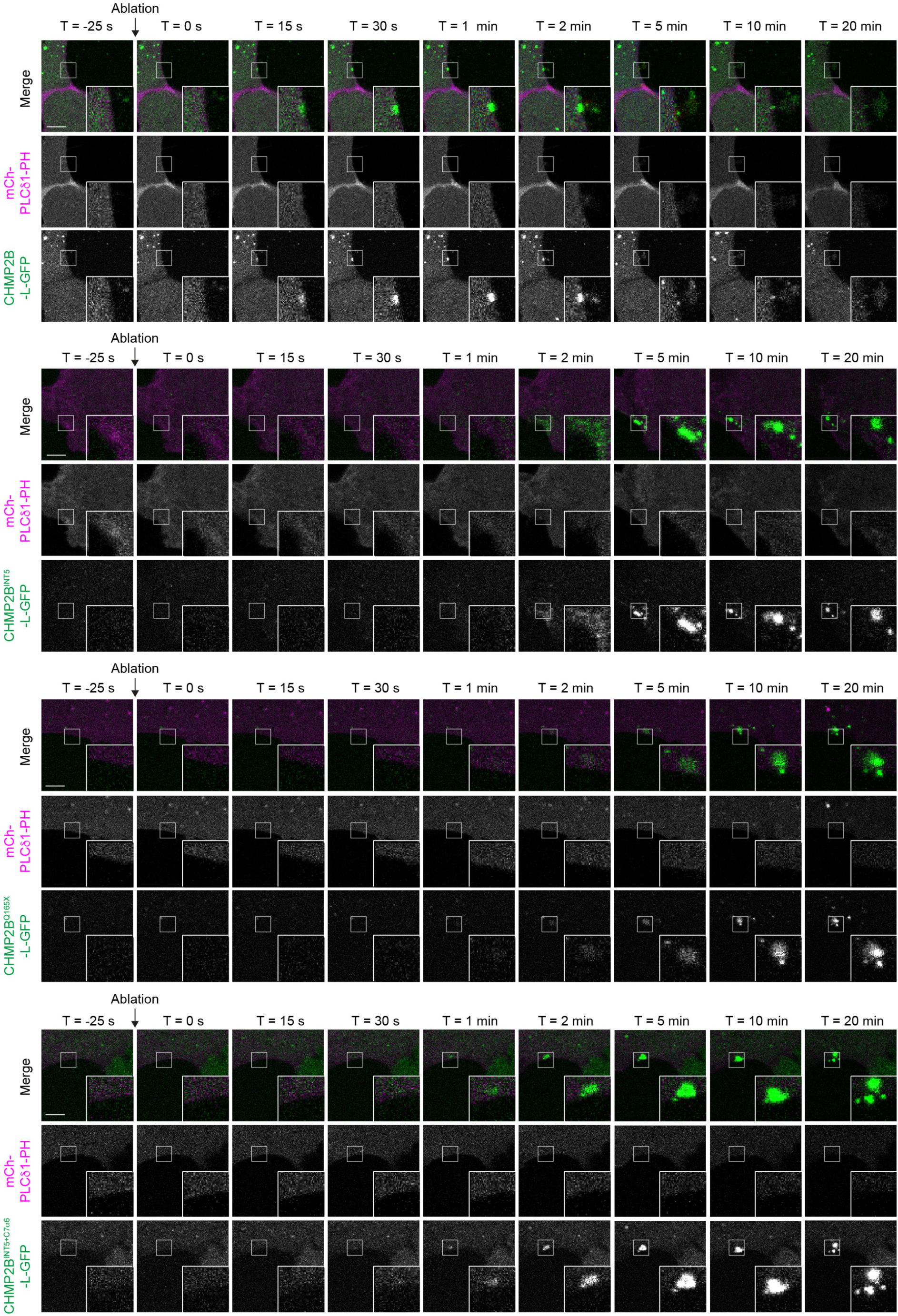
C-truncating variants of CHMP2B arrive late and persist at sites of plasma membrane damage. Extended sequence of representative images from cells described in Fig. 2A. CAL-51 cells stably expressing CHMP2B-L-GFP, CHMP2B^INT5^-L-GFP, CHMP2B^Q165X^-L-GFP or CHMP2B-L-GFP, CHMP2B^INT5+C7NES^-L-GFP were subject to a single-photon laser ablation at the mCh-PLC81-PH illuminated plasma membrane. Arrowheads indicate site of laser ablation at T = 0. In microscopy images, scale bar is 5 μm. See corresponding movies S1, S3-S5.

**Figure S6.**
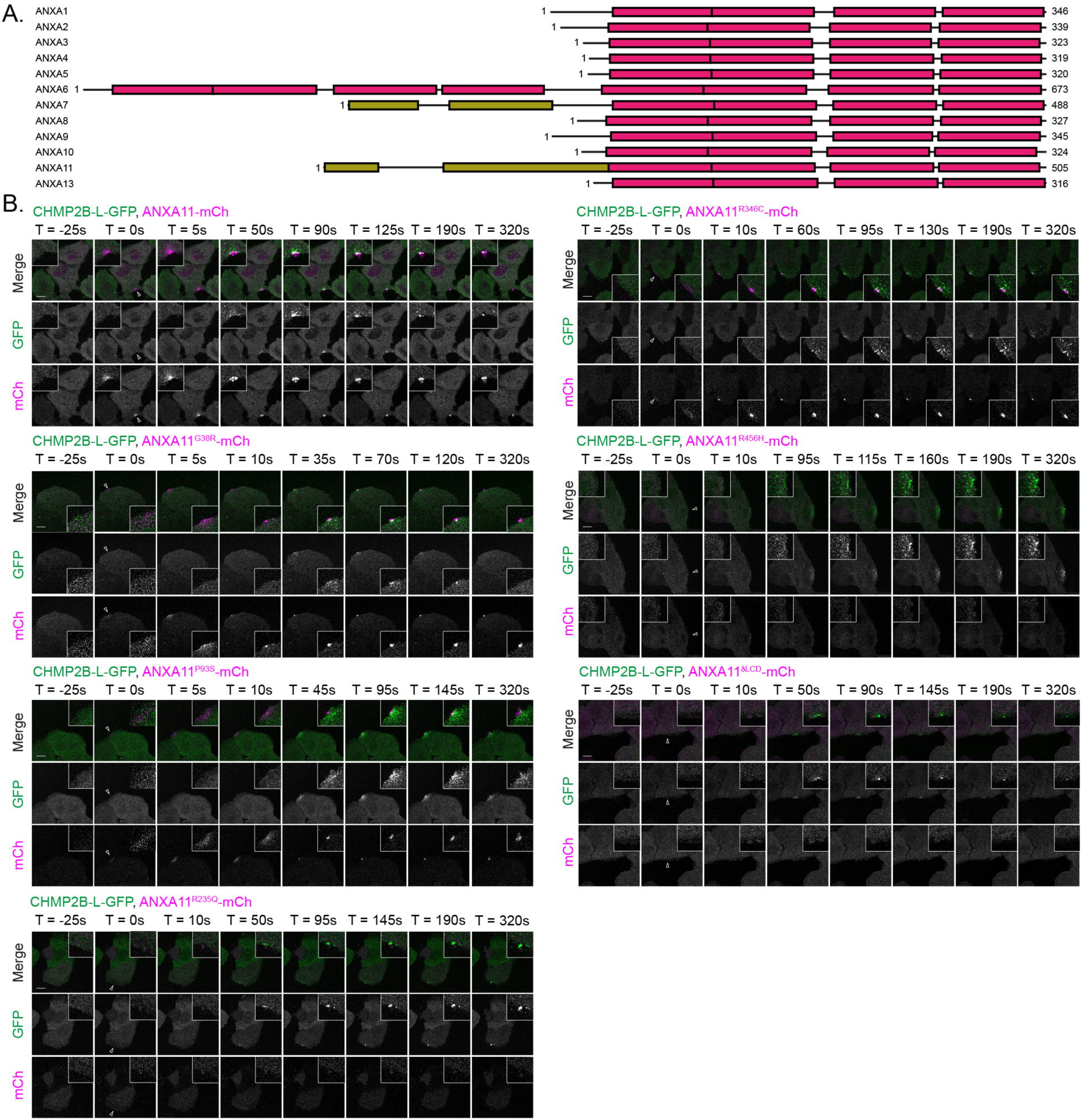
ALS-causing mutations in ANXA11 impair its assembly at sites of membrane damage. **A**. Schematic of sequences of human annexin proteins A1-A11. Annexin-like domain shown in pink, low-complexity domain shown in gold. **B**. Representative image sequences from cells described in Figure 5A. Representative images of CAL-51 cells stably expressing CHMP2B-L-GFP and either ANXA11-mCh, ANXA11^G38R^-mCh, ANXA11^P93S^-mCh, ANXA11^R235Q^-mCh, ANXA11^R456H^-mCh or ANXA11^8LCD^-mCh and subject to a single multi-photon laser ablation. Recruitment of CHMP2B-L-GFP (enlarged in boxed region) was used as a positive indicator of the membrane repair process. CHMP2B-L-GFP:ANXA11-mCh (n = 22, N = 8); CHMP2B-L-GFP:ANXA11^G38R^-mCh (n = 11, N = 5); CHMP2B-L-GFP:ANXA11^P93S^-mCh (n = 12, N = 4); CHMP2B-L-GFP:ANXA11^R235Q^-mCh, n = 10 cells, N = 4; CHMP2B-L-GFP:ANXA11^R346C^-mCh, n = 12 cells, N = 4; CHMP2B-L-GFP:ANXA11^R456H^-mCh, n = 9 cells, N = 2; CHMP2B-L-GFP:ANXA11^8LCD^-mCh, n = 15 cells, N = 5. In microscopy images, scale bar is 5 μm. See corresponding movies S10-S16.

**Figure S7.**
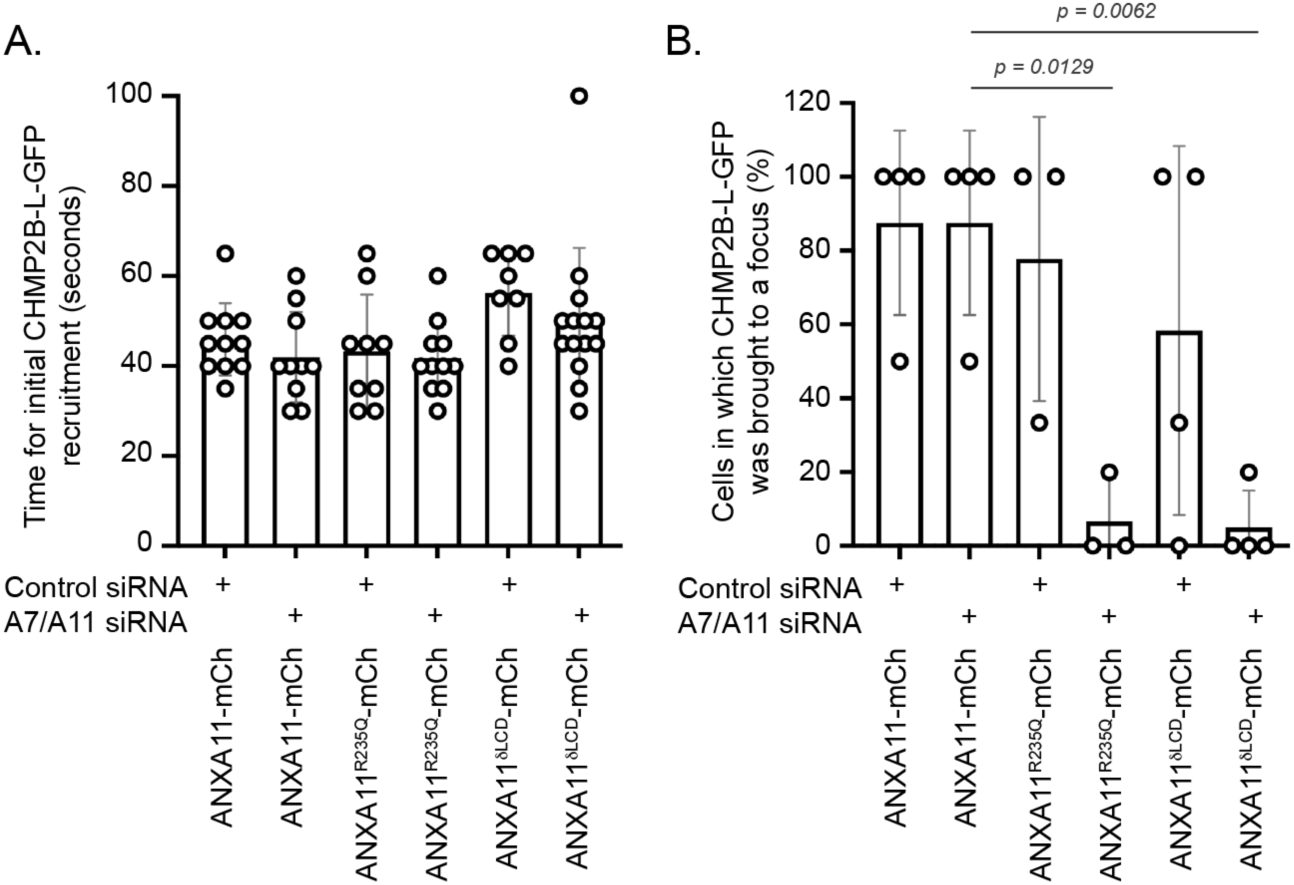
ANXA11 recruitment to sites of plasma membrane damage is needed for CHMPB-L-GFP turnover. **A**. Quantification of CHMP2B-L-GFP recruitment time in CAL-51 cells stably expressing CHMP2B-L-GFP, either ANXA11-mCh, ANXA11^R235Q^-mCh or ANXA11^8LCD^-mCh, transfected with control or ANXA7/ANXA11-targeting siRNA and subject to a single multi-photon laser ablation at the plasma membrane. Data acquired from N = 4 independent experiments, for which CHMP2B-L-GFP:ANXA11-mCh, control siRNA, n = 10; CHMP2B-L-GFP:ANXA11-mCh, ANXA7/ANXA11 siRNA, n = 10; CHMP2B-L-GFP:ANXA11^R235Q^-mCh, control siRNA, n = 9; CHMP2B-L-GFP:ANXA11^R235Q^-mCh, ANXA7/ANXA11 siRNA, n = 11; CHMP2B-L-GFP:ANXA11^8LCD^-mCh, control siRNA, n = 8; CHMP2B-L-GFP:ANXA11^8LCD^-mCh, ANXA7/ANXA11 siRNA, n = 15. Significance calculated by ordinary one-way ANOVA with Tukey’s correction. Significance was not achieved across any comparison. **B**. Quantification of whether the broad CHMP2B-L-GFP recruitment was brought to a focus at the site of membrane damage in A. Graph show mean ± S.E.M. CHMP2B-L-GFP:ANXA11-mCh, control siRNA, N = 4, n = 7; CHMP2B-L-GFP:ANXA11-mCh, ANXA7/ANXA11 siRNA, N = 4, n = 10; CHMP2B-L-GFP:ANXA11^R235Q^-mCh, control siRNA, N = 3, n = 7; CHMP2B-L-GFP:ANXA11^R235Q^-mCh, ANXA7/ANXA11 siRNA, N = 3, n = 8; CHMP2B-L-GFP:ANXA11^8LCD^-mCh, control siRNA, N = 4, n = 8; CHMP2B-L-GFP:ANXA11^8LCD^-mCh, ANXA7/ANXA11 siRNA, N = 4, n = 13. Significance calculated via ordinary one-way ANOVA with Dunnett’s correction.

**Figure S8.**
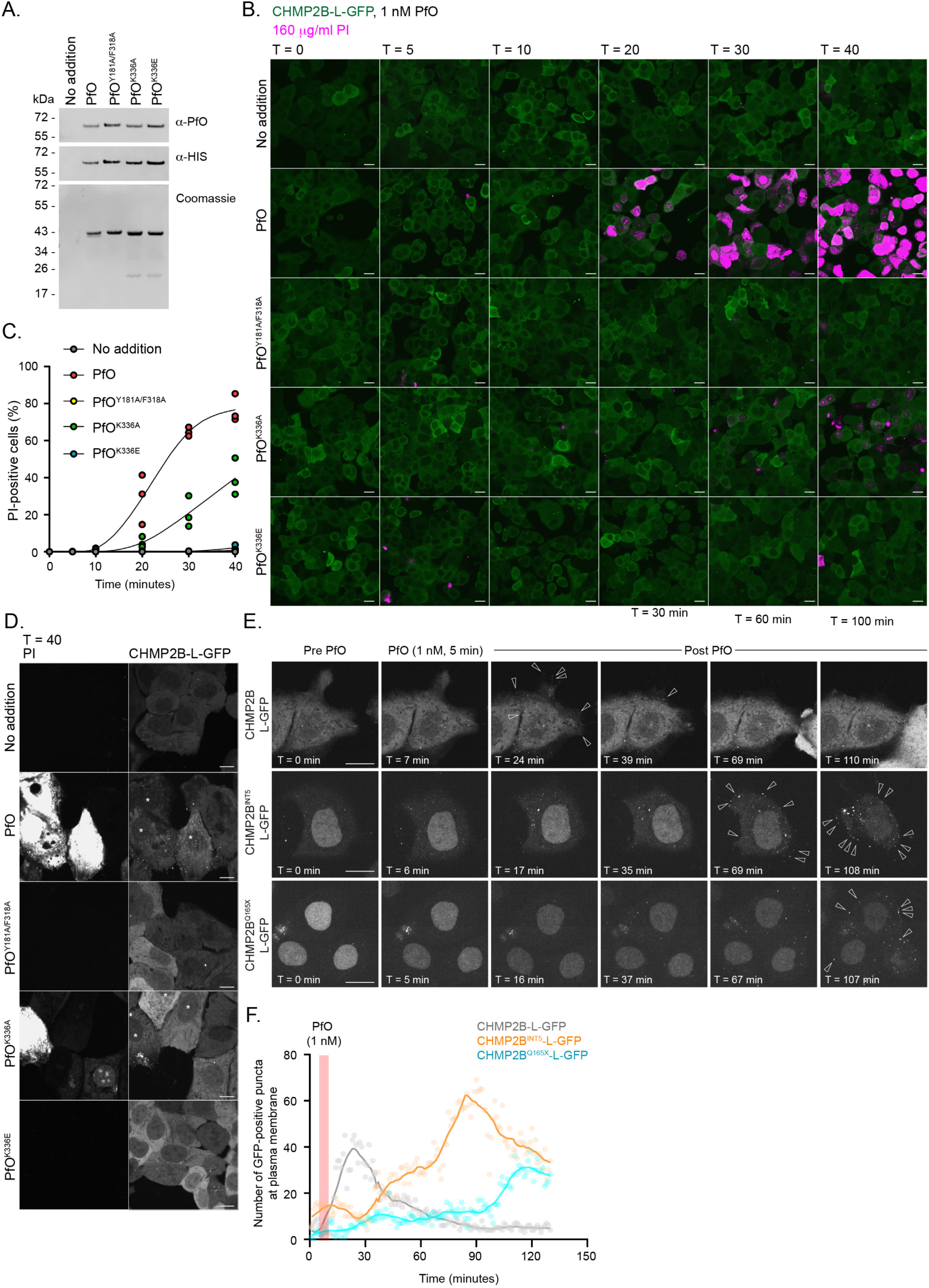
Characterisation of a non-lytic PfO and the response of C-truncated mutants of CHMP2B to PfO-mediated membrane damage. **A**. Resolved fractions of recombinant HIS-tagged PfO, PfO^Y181A/F316A^, PfO^K336A^ and PfO^K336E^ were examined by SDS-PAGE and western blotting with antisera raised against PfO, HIS or were examined by Coomassie staining. **B**. Representative image sequences of CAL-51 cells stably expressing CHMP2B-L-GFP and treated with vehicle or the indicated recombinant PfO proteins at 1 nM in the presence of 160 μg/mL PI and imaged live. Scale bar is 20 μM. **C**. Quantification of PI entry in cells described in B. Vehicle (n = between 746 and 1335 cells per timepoint; PfO (n = between 725 and 1276 cells per timepoint), PfO^Y181A/F316A^ (n = between 992 and 1403 cells per timepoint), PfO^K336A^ (n = between 931 and 1357 cells per timepoint) and PfO^K336E^ (n = between 1031 and 1382 cells per timepoint) from N = 3 independent experiments. **D**. Examination of CHMP2B-L-GFP puncta at the plasma membrane of cells from B. CHMP2B-L-GFP puncta are present in cells with PI influx (indicated by stars). Scale bar is 10 μM. **E, F**. Live cell imaging (E) and quantification (F) of CHMP2B puncta formation (indicated by arrowheads) at the plasma membrane of CAL-51 cells stably expressing CHMP2B-L-GFP, CHMP2B^INT5^-L-GFP or CHMP2B^Q165X^-L-GFP, imaged for 5 minutes (pre-PfO) treated with 1 nM PfO for 5 minutes and chased into fresh media (post-PfO). Scale bar is 10 μM. A Lowess curve was used to fit the data in C and F. See corresponding movies S28-S30.

**Figure S9.**
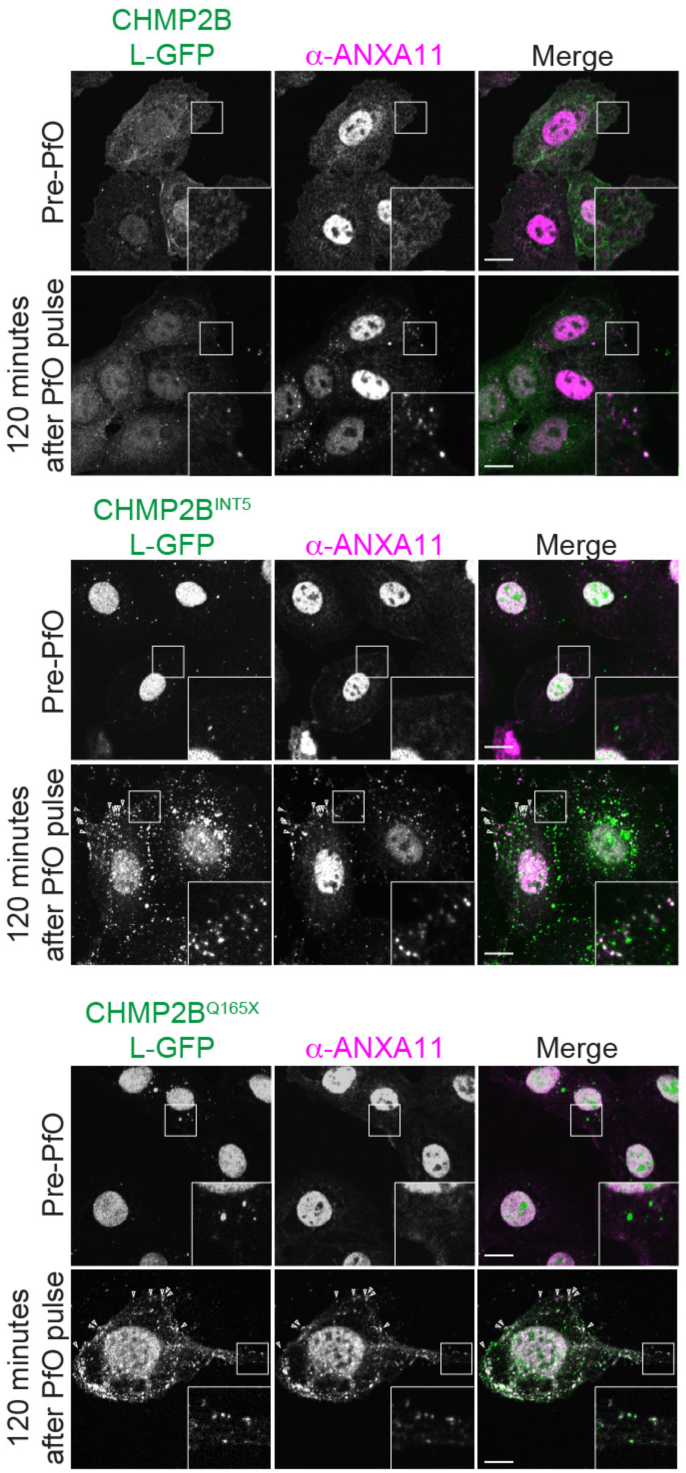
C-truncating mutations of CHMP2B cause retention of endogenous ANXA11 at sites of membrane damage. CAL-51 cells stably expressing CHMP2B-L-GFP, CHMP2B^INT5^-L-GFP or CHMP2B^Q165X^-L-GFP were treated with 1 nM PfO for 5 minutes, washed and chased into fresh media for 2 hours. Cells fixed and stained with antibodies raised against ANXA11. Cells displaying > 3 puncta at the plasma membrane where endogenous ANXA11 colocalised with the GFP was scored from N = 2 experiments (CHMP2B-L-GFP, 13± 3.1 % of cells, n = 92; CHMP2B^INT5^-L-GFP, 64 ± 5.3 % of cells, n = 77; CHMP2B^Q165X^-L-GFP, 100 %o f cells, n = 16)

## SUPPLEMENTAL MOVIE LEGENDS

**Movie S1: Recruitment of CHMP2B-L-GFP to sites of membrane damage.** CAL-51 cells stably expressing CHMP2B-L-GFP and transfected with mCh-PLC81-PH were imaged live and subject to a single multi-photon laser ablation at the plasma membrane (T = 0 seconds, site marked by arrowhead). Frames were acquired every 5 seconds, n = 14 movies, N = 7.

**Movie S2: Lamellipodia reveal wave-like kinetics of CHMP2B-L-GFP recruitment.** CAL-51 cells stably expressing CHMP2B-L-GFP and transfected with mCh-PLC81-PH were imaged live and subject to a single multi-photon laser ablation at the plasma membrane (T = 0 seconds, site marked by arrowhead). Frames were acquired every 5 seconds, n = 11 movies.

**Movie S3: Recruitment of CHMP2B^INT^**^5^**-L-GFP to sites of membrane damage.** CAL-51 cells stably expressing CHMP2B^INT5^-L-GFP and transfected with mCh-PLC81-PH were imaged live and subject to a single multi-photon laser ablation at the plasma membrane (T = 0 seconds, site marked by arrowhead). Frames were acquired every 5 seconds, n = 11 movies, N = 5.

**Movie S4: Recruitment of CHMP2B^Q^**^165^**^X^-L-GFP to sites of membrane damage.** CAL-51 cells stably expressing CHMP2B^Q165X^-L-GFP and transfected with mCh-PLC81-PH were imaged live and subject to a single multi-photon laser ablation at the plasma membrane (T = 0 seconds, site marked by arrowhead). Frames were acquired every 5 seconds, n = 12 movies, N = 6.

**Movie S5: Recruitment of CHMP2B^INT5+C^**^7^**^α^**^6^**-L-GFP to sites of membrane damage.** CAL-51 cells stably expressing CHMP2B^INT5+C7α6^-L-GFP and transfected with mCh-PLC81-PH were imaged live and subject to a single multi-photon laser ablation at the plasma membrane (T = 0 seconds, site marked by arrowhead). Frames were acquired every 5 seconds, n = 12 movies, N = 6.

**Movie S6: PI entry after laser ablation is transient and abates before CHMP2B-L-GFP is recruited.** CAL-51 cells stably expressing CHMP2B-L-GFP were imaged live in the presence of 160 μg/ml PI and subject to a single multi-photon laser ablation at the plasma membrane (T = 0 seconds, site marked by arrowhead). Frames were acquired every 5 seconds, n = 15 movies, N = 5.

**Movie S7: ALG2 is recruited to sites of membrane damage before CHMP2B.** CAL-51 cells stably expressing CHMP2B-L-GFP and ALG-2-mCh were imaged live and subject to a single multi-photon laser ablation at the plasma membrane (T = 0 seconds, site marked by arrowhead). Frames were acquired every 5 seconds, n = 5 movies, N = 2.

**Movie S8: ANXA7 is recruited to sites of membrane damage before CHMP2B.** CAL-51 cells stably expressing CHMP2B-L-GFP and ANXA7-mCh were imaged live and subject to a single multi-photon laser ablation at the plasma membrane (T = 0 seconds, site marked by arrowhead). Frames were acquired every 5 seconds, n = 8 movies, N = 3.

**Movie S9: ANXA11 is recruited to sites of membrane damage before CHMP2B.** CAL-51 cells stably expressing CHMP2B-L-GFP and ANXA11-mCh were imaged live and subject to a single multi-photon laser ablation at the plasma membrane (T = 0 seconds, site marked by arrowhead). Frames were acquired every 5 seconds, n = 11 movies, N = 3.

**Movie S10**: **ANXA11-mCh is recruited to sites of membrane damage in advance of CHMP2B-L-GFP.** CAL-51 cells stably expressing CHMP2B-L-GFP and ANXA11-mCh were imaged live and subject to a single multi-photon laser ablation at the plasma membrane (T = 0 seconds, site marked by arrowhead). Control experiments for mutants described in Movie S11 – S16. Frames were acquired every 5 seconds, n = 23 movies, N = 8.

**Movie S11: ANXA11^G38R^-mCh is recruited to sites of membrane damage in advance of CHMP2B-L-GFP.** CAL-51 cells stably expressing CHMP2B-L-GFP and ANXA11^G38R^-mCh were imaged live and subject to a single multi-photon laser ablation at the plasma membrane (T = 0 seconds, site marked by arrowhead). Frames were acquired every 5 seconds, n = 9 movies, N = 4.

**Movie S12: ANXA11^P93S^-mCh is recruited to sites of membrane damage in advance of CHMP2B-L-GFP.** CAL-51 cells stably expressing CHMP2B-L-GFP and ANXA11^P93S^-mCh were imaged live and subject to a single multi-photon laser ablation at the plasma membrane (T = 0 seconds, site marked by arrowhead). Frames were acquired every 5 seconds, n = 11 movies, N = 4.

**Movie S13: ANXA11^R^**^346^**^C^-mCh is recruited to sites of membrane damage in advance of CHMP2B-L-GFP.** CAL-51 cells stably expressing CHMP2B-L-GFP and ANXA11^R346C^-mCh were imaged live and subject to a single multi-photon laser ablation at the plasma membrane (T = 0 seconds, site marked by arrowhead). Frames were acquired every 5 seconds, n = 12 movies, N = 4.

**Movie S14: ANXA11^R^**^235^**^Q^-mCh is not recruited to sites of membrane damage.** CAL-51 cells stably expressing CHMP2B-L-GFP and ANXA11^R235Q^-mCh were imaged live and subject to a single multi-photon laser ablation at the plasma membrane (T = 0 seconds, site marked by arrowhead). Frames were acquired every 5 seconds, n = 12 movies, N = 5.

**Movie S15: ANXA11^R^**^456^**^H^-mCh is not recruited to sites of membrane damage.** CAL-51 cells stably expressing CHMP2B-L-GFP and ANXA11^R456H^-mCh were imaged live and subject to a single multi-photon laser ablation at the plasma membrane (T = 0 seconds, site marked by arrowhead). Frames were acquired every 5 seconds, n = 11 movies, N = 3.

**Movie S16: ANXA11^8LCD^-mCh is not recruited to sites of membrane damage.** CAL-51 cells stably expressing CHMP2B-L-GFP and ANXA11^8LCD^-mCh were imaged live and subject to a single multi-photon laser ablation at the plasma membrane (T = 0 seconds, site marked by arrowhead). Frames were acquired every 5 seconds, n = 16 movies, N = 6.

**Movie S17: Behavior of CHMP2B-L-GFP after membrane damage in ANXA7/ANXA11-depleted cells re-expressing siRNA resistant ANXA11-mCh.** CAL-51 cells stably expressing CHMP2B-L-GFP and siRNA resistant ANXA11-mCh were transfected with ANXA7 and ANXA11-targeting siRNA, imaged live and subject to a single multi-photon laser ablation at the plasma membrane (T = 0 seconds, site marked by arrowhead). Frames were acquired every 5 seconds, n = 10 movies, N = 4.

**Movie S18: Behavior of CHMP2B-L-GFP after membrane damage in ANXA7/ANXA11-depleted cells re-expressing siRNA resistant ANXA11^R^**^235^**^Q^-mCh.** CAL-51 cells stably expressing CHMP2B-L-GFP and siRNA resistant ANXA11^R235Q^-mCh were transfected with ANXA7 and ANXA11-targeting siRNA, imaged live and subject to a single multi-photon laser ablation at the plasma membrane (T = 0 seconds, site marked by arrowhead). Frames were acquired every 5 seconds, n = 11 movies, N = 4.

**Movie S19: Behavior of CHMP2B-L-GFP after membrane damage in ANXA7/ANXA11-depleted cells re-expressing siRNA resistant ANXA11^8LCD^-mCh.** CAL-51 cells stably expressing CHMP2B-L-GFP and siRNA resistant ANXA11^δLCD^-mCh were transfected with ANXA7 and ANXA11-targeting siRNA, imaged live and subject to a single multi-photon laser ablation at the plasma membrane (T = 0 seconds, site marked by arrowhead). Frames were acquired every 5 seconds, n = 14 movies, N = 4.

**Movie S20: PI entry at sites of PfO-mediated membrane damage is transient and CHMP2B-L-GFP is recruited after PI entry has abated.** CAL-51 cells stably expressing CHMP2B-L-GFP were treated with 1 nM PfO and imaged live in the presence of 160 μg/ml PI. Frames were acquired every 10 seconds and frames proximal to PI entry are presented.

**Movie S21: ANXA7-mCh and CHMP2B-L-GFP arrive sequentially at sites of PfO-mediated membrane damage.** CAL-51 cells stably expressing ANXA7-mCh and CHMP2B-L-GFP were treated with 1 nM PfO for 5 minutes, washed thrice, released into fresh imaging media and imaged live. Frames from N = 5 independently imaged cells were acquired every 10 seconds. Timestamp relative to appearance of ANXA7-mCh puncta.

**Movie S22: ANXA11-mCh and CHMP2B-L-GFP arrive sequentially at sites of PfO-mediated membrane damage.** CAL-51 cells stably expressing ANXA11-mCh and CHMP2B-L-GFP were treated with 1 nM PfO for 5 minutes, washed thrice, released into fresh imaging media and imaged live. Frames from N = 4 independently imaged cells were acquired every 10 seconds. Timestamp relative to appearance of ANXA11-mCh puncta.

**Movie S23: ANXA7-mCh puncta formed at the plasma membrane after PfO-mediated membrane damage are lost following recruitment of CHMP2B-L-GFP .** CAL-51 cells stably expressing ANXA7-mCh and CHMP2B-L-GFP were treated with 1 nM PfO for 5 minutes, washed thrice, released into fresh imaging media and imaged live. Frames from N = 3 independent experiments were acquired every 10 seconds. Timestamp relative to appearance of ANXA7-mCh puncta.

**Movie S24: ANXA11-mCh puncta formed at the plasma membrane after PfO-mediated membrane damage are lost following recruitment of CHMP2B-L-GFP.** CAL-51 cells stably expressing ANXA11-mCh and CHMP2B-L-GFP were treated with 1 nM PfO for 5 minutes, washed thrice, released into fresh imaging media and imaged live. Frames from N = 4 independent experiments were acquired every 10 seconds. Timestamp relative to appearance of ANXA11-mCh puncta.

**Movie S25: No response of actin cytoskeleton to laser ablation of the plasma membrane** CAL-51 cells stably expressing CHMP2B-L-GFP were transiently transfected with MP-Act-mRuby3 to illuminate the membrane proximal F-Actin. Cells were subjected to laser ablation of the plasma membrane and imaged live. Frames were acquired every 5 seconds.

**Movie S26: Disruption of actin cytoskeleton with cytochalasin D has no effect on CHMP2B-L-GFP recruitment to laser ablation damage at the plasma membrane** CAL-51 cells stably expressing CHMP2B-L-GFP were treated with 2uM cytochalasin D for 30 min, laser damaged at the plasma membrane, and imaged live. Frames were acquired every 5 seconds.

**Movie S27: No response of LAMP1 positive lysosomes to laser ablation of the plasma membrane** CAL-51 cells stably expressing CHMP2B-L-GFP were transiently transfected with LAMP1-tdTomato, laser damaged at the plasma membrane, and imaged live. Frames were acquired every 5 seconds.

**Movie S28: CHMP2B-L-GFP accumulates transiently at the plasma membrane of PfO-damaged cells.** CAL-51 cells stably expressing CHMP2B-L-GFP were treated with 1 nM PfO for 5 minutes, washed thrice, released into fresh imaging media and imaged live. Frames were acquired every minute.

**Movie S29: CHMP2B^INT^**^5^**-L-GFP accumulates persistently at the plasma membrane of PfO-damaged cells.** CAL-51 cells stably expressing CHMP2B^INT5^-L-GFP were treated with 1 nM PfO for 5 minutes, washed thrice, released into fresh imaging media and imaged live. Frames were acquired every minute.

**Movie S30: CHMP2B^Q^**^165^**^X^-L-GFP accumulates persistently at the plasma membrane of PfO-damaged cells.** CAL-51 cells stably expressing CHMP2B^Q165X^-L-GFP were treated with 1 nM PfO for 5 minutes, washed thrice, released into fresh imaging media and imaged live. Frames were acquired every minute.

